# Disentangling how multiple traits drive 2 strain frequencies in SIS dynamics with coinfection

**DOI:** 10.1101/2021.04.29.442023

**Authors:** Thi Minh Thao Le, Sten Madec, Erida Gjini

## Abstract

A general theory for competitive dynamics among many strains at the epidemiological level is required to understand polymorphisms in virulence, transmissibility, antibiotic resistance and other biological traits of infectious agents. Mathematical coinfection models have addressed specific systems, focusing on the criteria leading to stable coexistence or competitive exclusion, however, due to their complexity and nonlinearity, analytical solutions in coinfection models remain rare. Here we study a 2-strain SIS compartmental model with co-infection/co-colonization, incorporating multiple fitness dimensions under the same framework: variation in transmissibility, duration of carriage, pairwise susceptibilities to coinfection, coinfection duration, and transmission priority effects from mixed coinfection. Taking advantage of a singular perturbation approach, under the assumption of strain similarity, we expose how strain dynamics on a slow timescale are explicitly governed by a replicator equation which encapsulates all traits and their interplay. This allows us to predict explicitly not only the final epidemiological outcome of a given 2-player competition, but moreover, their entire frequency dynamics as a direct function of their relative variation and of strain-transcending global parameters. Based on mutual invasion fitnesses, we analyze and report rigorous results on transition phenomena in the 2-strain system, strongly mediated via coinfection prevalence. We show that coinfection is not always a promoter of coexistence; instead, its effect to favour or prevent polymorphism is non-monotonic and depends on the type and level of phenotypic differentiation between strains. This framework offers a deeper analytical understanding of 2-strain competitive games in coinfection, with theoretical and practical applications in epidemiology, ecology and evolution.

## 1 Introduction

Epidemiological models of coinfection have a long history of study (Levin and Pimentel, 1981; Adler and Brunet, 1991; Nowak and May, 1994; May and Nowak, 1995; van Baalen and Sabelis, 1995; Mosquera and Adler, 1998; Martcheva, 2009; Thieme, 2007; Alizon, 2013). Examples of multi-strain infectious agents where coinfection processes appear and shape epidemiology include *Streptococcus pneumoniae* bacteria (Lipsitch, 1997; Gjini et al., 2016), *Bordetella pertussis* (Nicoli et al., 2015), *Mycobacterium tuberculosis* (Cohen et al., 2012), *Staphylococcus Aureus*, (Pinotti et al., 2019) and many others, comprising plants (Susi et al., 2015; Halliday et al., 2020), and also inter-species co-colonization such as between *Haemophilus Influenzae* and pneumococcus serotypes (Margolis et al., 2010; Cobey and Lipsitch, 2013) and coinfection with different viruses (Furuya-Kanamori et al., 2016). Typically the strain-defining parameters vary much less within than between species. However, until now, models have not leveraged the conceptual and analytic advantages of strain similarity to the full extent, except for the classical comparisons between strain-specific basic reproduction numbers.

While it has long been recognized that in coinfection systems, basic reproduction numbers alone do not determine strain competitive dynamics (Nowak and May, 1994; van Baalen and Sabelis, 1995), a generic framework to integrate variation among strains along several phenotypic axes and coinfection, and map these directly to strain frequencies, has not been developed. Moreover, until now simplified versions of traits involved in SIS dynamics between two strains have been addressed: either modeling vulnerability to coinfection as a single parameter (Alizon, 2013), or focusing just on cross-strain competition (Lipsitch, 1997), four-way competitive interactions via altered susceptibilities to coinfection (Gjini et al., 2016), exclusively cooperative dynamics (Chen et al., 2017), or focusing on transmission and clearance rate variation (Martcheva, 2009; Thieme, 2007).

A key level of strain interactions in coinfection is the within-host level, where the order and timing of arrival, can matter for onward transmission or clearance. Such interactions when studied empirically have revealed strong priority effects, where the first arriving genotype has an advantage over later arriving ones (De Roode et al., 2005; Halliday et al., 2020). Independently of the underlying mechanisms, whether via host immunity, resource overlap within host or others, priority effects have repercussions on disease dynamics and parasite assemblage dynamics at higher scales. Yet, the full extent of the inter-dependence between this trait and other traits involved in epidemiological dynamics remains poorly understood. Thus, although several aspects of coinfection have been studied, typically with simulations or analyses restricted to special cases and particular models, a comprehensive and concise theoretical framework for how coinfection prevalence broadly interplays with multiple traits between 2 coinfecting agents in endemic systems is still missing.

In this paper, we describe and study a general system for epidemiological dynamics of similar co-infecting entities (e.g. discrete strains of the same infectious agent or similar species) that comprise a rich ecological and epidemiological phenomenology. Such a system could apply, but not be limited to, polymorphic *Streptococcus pneumonia* bacteria or other commensal bacteria (Lipsitch, 1997; Cobey and Lipsitch, 2012; Gjini et al., 2016; Davies et al., 2019). We generalize a previously-introduced quasi-neutral *Susceptible-Infected-Susceptible* (SIS) framework for 2 circulating strains and co-infection, where we showed that asymmetries in pairwise susceptibilities to co-infection create frequency-dependent advantage for one of the strains and can give rise to coexistence, bistability as well as competitive exclusion between strains (Gjini et al., 2016; Gjini and Madec, 2017). Here, we study additional variation between two microbial strains, namely, in other traits besides vulnerability to co-infection, including transmission rate and duration of carriage, two classical traits that are known to vary among colonizing pneumococcus serotypes (Abdullahi et al., 2012), but in general can also vary among two arbitrary infectious agents, e.g. an antibiotic-resistant and an antibiotic-sensitive strain. Furthermore, we also allow for variation in transmission biases from co-infected hosts carrying a mixture of two strains, and in duration of coinfection episodes, where priority effects can play a role, adding new layers where competitive abilities and asymmetries can manifest. Until now analytic solutions for how such systems behave in time have not been obtained, although theoretical models have studied the conditions for coexistence vs. competitive exclusion or used numerical simulations as an approach in specific cases (Lipsitch, 1997; Zhang et al., 2004; Gjini et al., 2016; Alizon, 2008; Hansen and Day, 2014).

The novelty of our approach lies in applying singular perturbation theory to a quasi-neutral model, whereby we obtain a timescale separation, similar to (Gjini and Madec, 2017), in order to express the total dynamics as a fast plus a slow component, related to broken symmetries along 5 traits between strains: transmission rate *β*_*i*_, clearance rate *γ*_*i*_, coinfection duration *γ*_*ij*_, pairwise vulnerabilitites to coinfection *k*_*ij*_, and transmission priority effects from coinfection 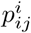. Net competition dynamics can be complex because all traits interact nonlinearly to determine final strain fitness at the host population level, but here we make such selective process entirely explicit. Moreover, we show how the strain-transcending parameters, defining the neutral model at the center, feed back on the strain dynamics on the slow timescale, and tune the net importance of each phenotypic axis.

The bigger *N* -strain SIS model with coinfection is derived in (Le et al., 2021), where all the formal mathematical details leading to a replicator equation that governs multi-strain frequency dynamics are provided. Here, we apply this framework to the simpler *N* = 2 case, and study in depth the emergent ecology mediated by different epidemiological traits. While in the general *N* − strain case analytical expressions for equilibrium frequencies can only be obtained in special cases, as shown by (Madec and Gjini, 2020; Le et al., 2021), in the *N* = 2 system, considered here, equilibrium frequencies are entirely explicit. This allows us to provide exact analytical results on qualitative and quantitative shifts in system behavior (coexistence vs. exclusion) with coinfection prevalence across a wide spectrum of biological scenarios.

As in Gjini and Madec (2017); Madec and Gjini (2020), we assume only up to 2 strains may co-infect a host (MOI=2). We model how primary infection by one strain alters host susceptibility to secondary strains, (increasing or decreasing it) by a factor *k*_*ij*_, relative to uninfected hosts, without acquired immunity. The altered susceptibilities to co-infection, given by a 2×2 matrix in the case of 2 strains, can comprise antagonistic or facilitative interactions (*k*_*ij*_ < 1 or ≥ 1). Beyond enabling competition and cooperation to be studied under the same framework, our model allows also for any asymmetries in this coinfection susceptibility matrix, as for the coinfection clearance rate matrix, depending on strain composition, and for transmission biases from coinfected hosts, depending on order of strain arrival.

Considering the complex epidemiological multi-strain dynamics in fast and slow components has many analytic and computational advantages as argued in (Madec and Gjini, 2020; Le et al., 2021). Our neutral model satisfies the criteria for ecological and population-genetic neutrality discussed in the context of ‘no coexistence for free’ (Lipsitch et al., 2009), but much more than a neutral null model, our approach highlights the neutral model as the core organizing centre of the multi-strain dynamics. This is made entirely explicit via the slow-fast timescale separation formalism (Madec and Gjini, 2020; Le et al., 2021), linking neutral and non-neutral dynamics in an ‘organic’ manner, and showing how parameters of the neutral model impact significantly on the slow frequency-dependent selection dynamics between strains, and tune the ecological feedbacks among different traits.

The paper is organized as follows. First, we describe the epidemiological framework. Secondly, we expose and elaborate on a closed and generic analytic solution for 2-strain frequency dynamics over the slow time-scale, in a changing fitness landscape shaped by multi-trait variation. This solution coincides with a version of the classical replicator equation in 2 dimensions (Hofbauer and Sigmund, 2003), but with an explicit payoff matrix derived from 5-dimensional trait variation between strains relatively weighted in the overall pairwise invasion fitness (Le et al., 2021). Third, we analyze why and how coexistence, bistability or competitive exclusion of either strain may occur between any two strains, for a fixed given trait variation between them. Fourth, we focus our attention on an in-depth analytic investigation of how strain-transcending mean-field gradients can shift the same system across these regimes, for different values of global *R*_0_ or coinfection prevalence, detailing the context-dependence of net outcomes. Finally, we conclude with a roadmap for biological applications. We believe our analysis and approach offer a fresh perspective, to quantify and predict how multiple traits together shape coinfecting strain dynamics and final equilibria via joint and nonlinear population feedbacks.

## 2 The modeling framework

### 2.1 The SIS model with coinfection

We study an infectious agent transmitted in a host population following Susceptible-Infected-Susceptible dynamics, where there are two co-circulating strains (denoted by 1 and 2). Susceptible hosts *S* can acquire any strain *i*, by which they enter the single infection (colonization) compartment *I*_*i*_. Singly-infected hosts *I*_*i*_ can acquire any secondary strain *j*, leading them to enter the coinfection (co-colonization) compartment *I*_*ij*_. As in the pioneering model by (van Baalen and Sabelis, 1995), an important epidemiological feature here is that hosts can be coinfected twice by the same strain (*I*_11_ and *I*_22_ compartments). Without this assumption, a rare strain always has an advantage: it can infect hosts already infected by the common strain while the common strain has few hosts to coinfect. Co-colonized and singly-colonized hosts transmit at equal total rate, and hosts carrying a mixture of two different strains transmit any strain *i* with a given probability 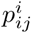 which can be different from 1/2 and may depend on the order of arrival within-host 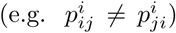. The model follows the structure in (Gjini et al., 2016; Gjini and Madec, 2017), but here we have a more general model, allowing for more trait variation between strains. In (Gjini and Madec, 2017), only pairwise susceptibilities to co-infection were modeled as different between strains (2-by-2 matrix of *k*_*ij*_ coefficients), and this was sufficient to generate stabilizing mechanisms for coexistence. Here, in addition to *k*_*ij*_, we model strain-specific transmission rate, *β*_*i*_, and clearance rate *γ*_*i*_, as well as coinfection clearance rates *γ*_*ij*_ and transmission biases from coinfection 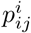, depending on strain composition in coinfection. Thus we describe two types of priority effects: at the between-host level in terms of the *k*_*ij*_, and at the within-host level in terms of the 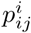. Recruitment of susceptibles happens at per-capita rate *r*, assumed equal to the natural mortality rate. The scheme of the model for two strains is given in Figure 1. The explicit dynamical system of equations for the *N* -strain version of this epidemiological model is derived in (Le et al., 2021), and given by:

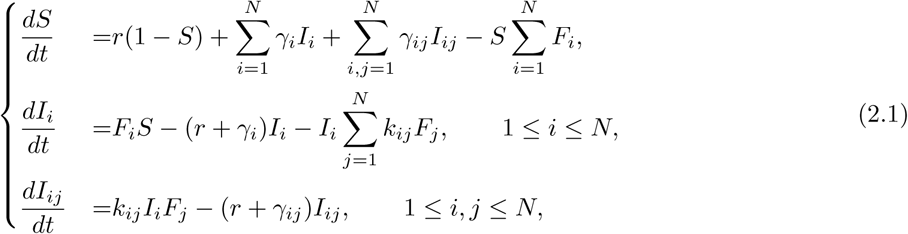

where in our case for *N* = 2, *i, j* ∈ {1, 2}, and the force of infection for each strain is given by:

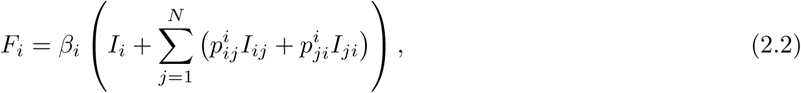

**Figure 1:**
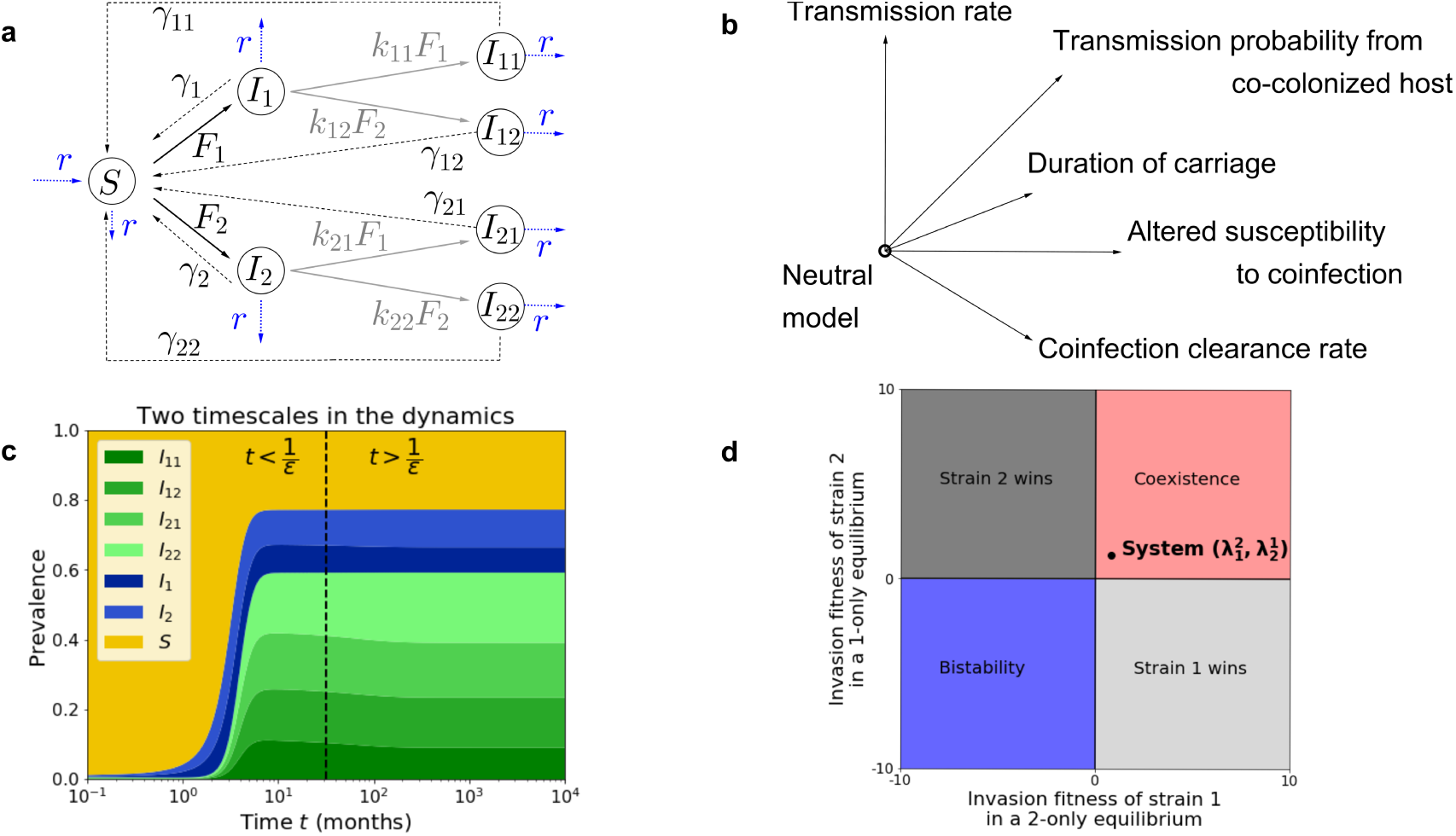
SIS coinfection model diagram for two strains (*N* = 2) and multivariate selection dynamics. **a**. The model follows the structure in (Gjini and Madec, 2017) but here two strains can differ in: transmission rate *β*_*i*_, clearance rate *γ*_*i*_, co-colonization clearance rate *γ*_*ij*_, altered susceptibilities to co-colonization *k*_*ij*_ and transmission biases from coinfection 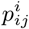. Thus any combination of relative fitness costs and advantages can be encapsulated, provided that their variation is not too big, as expected for similar conspecific strains, or similar infectious co-circulating ‘species’. Non-carriers (*S*) become carriers of either strain 1 or 2 (*I*_*i*_) with force of infection 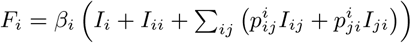 where the mixed carriage compartment (*I*_*ij*_) may transmit either strain with a slightly biased probability away from 1/2 depending on the order of arrival (see (Le et al., 2021)). Here 1*/γ*_*i*_ is the strain-specific duration of single colonization, 1*/γ*_*ij*_ are the composition-specific durations of co-colonization, which can vary for all four *I*_*ij*_ classes. The coefficients *k*_*ij*_ capture the altered relative susceptibilities to co-colonization between strains, when a host is already colonized, and transitions from primary colonization to co-colonization. The parameter *r* is the natural birth/death rate of the host. **b**. Assuming strain similarity, the epidemiological dynamics in such an SIS model with coinfection, can be decomposed into a fast (neutral) component and slow (non-neutral) component. The slow dynamics are shown to follow an explicit replicator equation which includes in the net payoff matrix variation across 5 dimensions of fitness for each strain (Le et al., 2021). This equation allows to predict analytically the entire temporal dynamics of two strains as a function of their epidemiological phenotypes. **c**. We simulate an example of 2-strain system in two timescales. On the fast time-scale (*o* (1*/ϵ*)), strains follow neutral dynamics, driven by mean-field parameters, where total prevalence of susceptibles, single infection and co-infection stabilize. On a slow time-scale, *ϵt*, within conserved global epidemiological compartments, complex non-neutral dynamics between strains takes place, depicted here by the blue and green shadings. **d**. Each system can be in one of four scenarios between 2 strains, depending on the signs of mutual invasion fitnesses (e.g. dynamics in **c** corresponds to the black point in the coexistence region). We find that frequency dynamics are explicitly governed by the 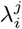. In our model, invasion fitnesses are explicit functions of strain variability along different traits and global mean-field parameters.

### 2.2 The similarity assumption

Some conventions and notations for the parameters in a multiple-trait model, under the similarity assumption between strains, are given in Table 1, where strain-specific parameters are defined in terms of relative variation from a common reference. For example, strain-specific transmission rates are described as *β*_*i*_ = *β*(1 + *ϵb*_*i*_). In particular, in this approach *ϵ* gives the scale of the perturbation around neutrality, which must be relatively small for the quasi-neutral approximation to hold, whereas the magnitudes of the perturbations in different traits are captured by Δ*b* = *b*_*i*_ − *b*_*j*_, etc. which may be large. Furthermore, it is important to note that there is not a unique representation in terms of ‘*scale* ×*magnitude*’ of perturbations for a particular system. For example there are many possibilities preserving 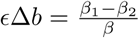, for fixed *β*_1_, *β*_2_ close to each other. In our framework, the magnitudes and directions of the perturbations in transmission rates, given by *b*_1_, *b*_2_ provide a description of how far proportionally from the common reference *β* are the two respective transmission rates, when measured in the scale of *ϵ*. Since we have many traits varying simultaneously, the choice of the appropriate *ϵ* must be convenient in order to describe all parameters under a common scale. In practice, capturing multivariate dissimilarity between two strains under the same scale, will involve an intermediate choice of *ϵ* balancing the requirement of relatively small for the approximation to hold, and sufficient for the slow dynamics to be of significant ecological/observational relevance. However, mathematically speaking, for the slow-fast method to be applied, the only requirement is that *ϵ* has to be as small as needed and that the perturbations *b*_*i*_, *ν*_*i*_ etc. are fixed.

**Table 1:** Conventions and notations of parameters and variables, where we assume the traits are numerically close for closely-related strains, or similar infectious entities. This similarity assumption (0 < *ϵ* ≪ 1, small) forms the basis for dynamic decomposition into fast and slow components (Le et al., 2021).

| Parameter | Interpretation | Specification | Features |
| --- | --- | --- | --- |
| **Original system (Quasi-neutral)** |  |  |  |
| $\beta_i = \beta(1 + \epsilon b_i)$ | Strain-specific transmission rates | $\Delta b = b_1 - b_2$ | Favours 1 if $\Delta b > 0$ |
| $\gamma_i = \gamma(1 + \epsilon \nu_i)$ | Strain-specific clearance rates of single colonization | $\Delta \nu = \nu_2 - \nu_1$ | Favours 1 if $\Delta \nu > 0$ |
| $\gamma_{ij} = \gamma(1 + \epsilon u_{ij})$ | Clearance rates of co-colonization with $i$ and $j$ | $\Delta_i u = 2u_{jj} - (u_{ij} + u_{ji})$ | Favours $i$ if $\Delta_i u > 0$ |
| $p_{ij}^s = \frac{1}{2} + \epsilon \omega_{ij}^s$ | Transmission probability <sup>1</sup> of $s \in \{i, j\}$; from a host co-colonized by strain-$i$ then -$j$ (priority effects). | $\Delta \omega = \omega_{12}^1 - \omega_{21}^2$ | Favours 1 if $\Delta \omega > 0$ |
| $k_{ij} = k + \epsilon \alpha_{ij}$ | Relative factor of altered susceptibility to co-colonization between colonizing strain $i$ and co-colonizing strain $j$ | $\Delta_i \alpha = \alpha_{ji} - \alpha_{jj} + \mu(\alpha_{ji} - \alpha_{ij})$ | Favours $i$ if $\Delta_i \alpha > 0$ |
| **Embedded neutral system (Fast):** |  |  |  |
| $r$ | Susceptible recruitment rate | Equal to natural mortality | Host turnover ↑ |
| $\beta$ | Transmission rate (infectiousness) | $\beta > 0$ | Transmission ↑ |
| $\gamma$ | Clearance rate of single infection | Equal to clearance rate of co-infection, $\gamma > 0$ | Transmission ↓ |
| $m$ | Net infected host turnover rate | $m = r + \gamma$ | $R_0 \downarrow$ |
| $R_0$ | Basic reproduction number | $R_0 = \beta/m > 1$ | Colonization ↑ |
| $k$ | Altered susceptibility to co-infection when infected | $k > 0$ | $k \geq 1$: facilitation,<br>$k < 1$: competition |
| $\mu$ | Single to co-infection ratio | $\mu = \frac{1}{(R_0 - 1)k}$ | Monotone in $R_0$ and $k$ |
| **Non-neutral system (Slow):** |  |  |  |
| $\theta_1$ | Weight of transmission rate axis $\beta_i$ | $(\mu + 1)^2$ | Depends on $\mu$ |
| $\theta_2$ | Weight of clearance rate axis $\gamma_i$ | $\frac{\gamma}{2(\gamma+r)}(2\mu^2 + \mu)$ | Depends on $\mu, \gamma, r$ |
| $\theta_3$ | Weight of co-infection clearance rate axis $\gamma_{ij}$ | $\frac{\gamma}{2(\gamma+r)}(\mu + 1)$ | Depends on $\mu, \gamma, r$ |
| $\theta_4$ | Weight of transmission priority effects from co-infection axis $p_{ij}^i$ | $\mu + 1$ | Depends on $\mu$ |
| $\theta_5$ | Weight of susceptibilities to co-infection axis $k_{ij}$ | $\frac{1}{2k}$ | Depends on $k$ |

To obtain the fast-slow decomposition, we rewrite the system 2.1 in terms of new aggregate variables as in (Madec and Gjini, 2020), such as the total prevalence of colonized hosts *T*, the total prevalence of hosts transmitting either strain *J*_*i*_, the total prevalence of single colonization *I*, and co-colonization *D*:

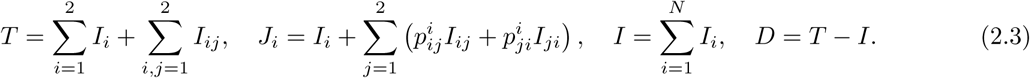

Following similar technical steps as in (Madec and Gjini, 2020), and applying Tikhonov’s theorem (Tikhonov, 1952), we have derived for the *N* strain model in (Le et al., 2021) that during the fast timescale, strains behave as neutral (all parameters are identical between them) and each global aggregated variable tends to its equilibrium: *S* → *S*^∗^, *T* → *T* ^∗^, *I* → *I*^∗^ and *D* → *D*^∗^ as *ϵ* → 0. With the basic reproduction number in this system denoted 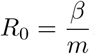, this equilibrium is given by:

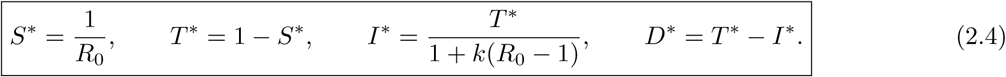

and the ratio of single infection to co-infection is given by 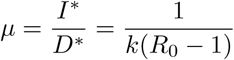.

Further, during the slow time scale *ϵt*, strains are not equivalent, their differences in fitness start to get manifested, and what follows is non-neutral dynamics at the level of strain frequencies *z*_*i*_. For a 2-strain system, there are four possibilities for the equilibrium: i) coexistence, ii-iii) exclusion of each strain, and iv) bistability of competitive exclusion states, also known as a priority effect. Below, these outcomes and their dynamics are shown to depend explicitly on mutual invasion growth rates between two strains.

### 2.3 Pairwise invasion fitness and replicator dynamics via timescale separation

In this more complex model, we follow the same reasoning as in (Madec and Gjini, 2020), focusing on mutual invasion fitnesses, to express the selective dynamics occurring on the slow time scale. We will define 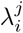 to be invasion fitness of strain *i* in an equilibrium set by strain *j* alone, a classical approach in adaptive dynamics (Geritz et al., 1998). Initially, based on (Le et al., 2021), we redefine *θ*_*i*_ via the global quantities and parameters of the neutral model:

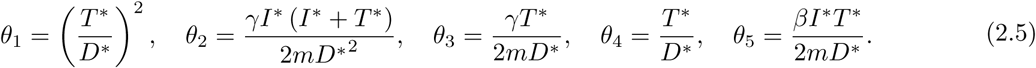

As derived in detail in (Le et al., 2021), we have that, in our model with multi-trait variation between strains, for *i, j* ∈ {1, 2}, the mutual invasion fitnesses are given by:

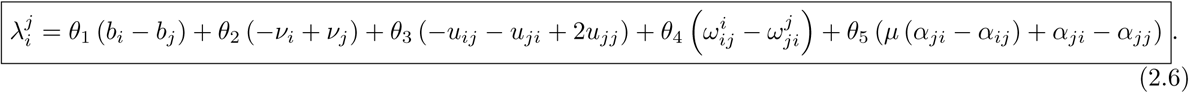

This analytic expression sums the relative contributions of multiple trait variations at the same time, with the weighting constants *θ*_*i*_ defined above and given explicitly in Table 1. By the notations of 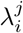, setting 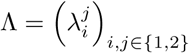, system (2.1) on the slow timescale *ϵt*, can be approximated by the replicator equation

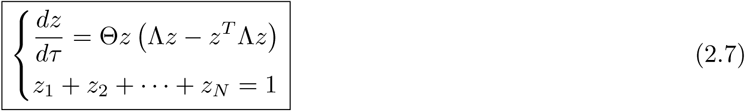

for variables *z* = (*z*_1_, *z*_2_) denoting strain frequencies, where the overall speed of dynamics Θ is given by:

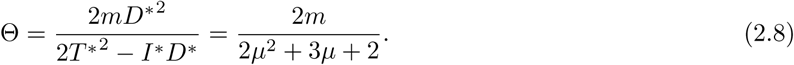

When there is just variation in co-infection susceptibility coefficients *k*_*ij*_, we recover the model and the Θ in (Madec and Gjini, 2020). Further, recall in (Madec and Gjini, 2020) the term *z*^*T*^ Λ*z* is denoted as *Q* and referred to as mean invasibility of the system, capturing resistance of the system to invasion by outsiders (Gjini and Madec, 2021b). In the 2-strain system, more explicitly we have: 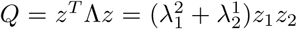, and this quantity is positive only in the case of coexistence between two strains.

In this slow-fast derivation, epidemiological variables of the original model (system 2.1) are then a function of strain frequencies of the slow system:

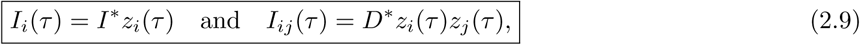

where *I*^∗^ and *D*^∗^ give the overall prevalence of single infection and co-infection in the endemic system (neutral model), and *z*_*i*_ and *z*_*i*_*z*_*j*_ give the proportions occupied by strain *i* and the pair of strains *i* and *j*, among singly infected hosts and co-infected hosts respectively.

## 3 General outcomes of the 2-strain system

### 3.1 Equilibria of the system

Denote by 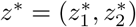 the nonzero equilibrium state of (2.7), where strain frequencies are given by:

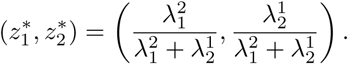

Depending on the signs of both invasion growth rates, we therefore have conditions for 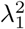 and 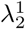, leading to four ecological scenarios between two strains as in (Madec and Gjini, 2020) (see Table 2). Thus, to investigate the equilibria and their stability in this 2-strain system, it suffices to study the values and signs of pairwise invasion fitness coefficients 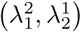, given explicitly as follows:

**Table 2:** System equilibria for 2-strain dynamics according to 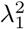 and 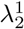 as expected from the replicator equation 2.7

| Mutual invasion<br>($\lambda_1^2, \lambda_2^1$) | Outcome | Strain frequencies | Quadratic form<br>$Q = z^T \Lambda z$ |
| --- | --- | --- | --- |
| (+, +) | Stable coexistence | $z_1^* > 0, z_2^* > 0$: $\frac{1}{\lambda_1^2 + \lambda_2^1} \begin{pmatrix} \lambda_1^2 \\ \lambda_2^1 \end{pmatrix}$ | $Q > 0$ and $Q \rightarrow Q^*$ |
| (-, +) | Exclusion of type 1 | $z_1^* = 0, z_2^* = 1$ | $Q \rightarrow 0$ |
| (+, -) | Exclusion of type 2 | $z_1^* = 1, z_2^* = 0$ | $Q \rightarrow 0$ |
| (-, -) | Bistability | $z_1^* = 1, \text{or } z_2^* = 1$ | $Q < 0$ and $Q \rightarrow 0$ |

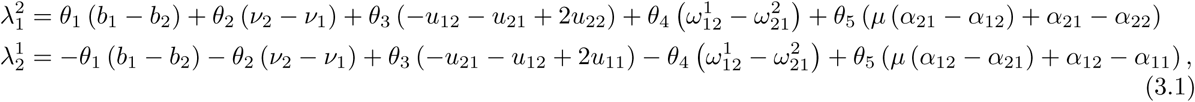

while their sum is 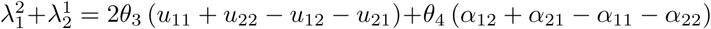. It’s not easy to determine the exact long time scenario or the winner in a two-strain system because parameters with perturbations affect all together the dynamics. The table 2 gives us criteria to determine the long time behavior. However, instead of computing the fitness coefficients explicitly in each case, we can base on (3.1) to determine quickly and for a various range of cases. Inspecting closely equations (3.1), we can see two parts:

1. The part *θ*_1_Δ*b* + *θ*_2_Δ*ν* + *θ*_4_Δ*ω* + *θ*_5_*µ* (*α*_21_ − *α*_12_), which keeps the 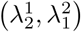 close to the line 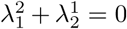, i.e. the dynamics tend to exclusion of one strain.
2. The part *θ*_3_Δ_2_*u* + *θ*_5_ (*α*_21_ − *α*_22_), which pulls 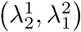 away from the line 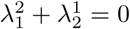, i.e. driving the dynamics toward coexistence or bi-stability.

This makes it easy to see that variation in transmissibility (Δ*b*) or duration of infection between strains (Δ*ν*), and the precedence effect in transmission from mixed coinfection (Δ*ω*), always promotes competitive exclusion in the system, whereas variation in coinfection parameters (susceptibilities and clearance rates) can oppose competitive exclusion.

### 3.2 An overview on four system outcomes dependent on *R*_0_ and *k*

#### What determines competitive exclusion?

The competitive exclusion occurs if and only if 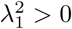 and 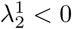 or reversely, 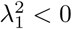 and 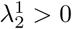. By (3.1), taking *R*_0_ → 1^+^ or *k* → 0^+^, which implies *µ* → ∞, makes the nonlinear part tends to 0, which leads to the competitive exclusion. This remark coincides with the result about 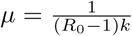 in (Gjini and Madec, 2021a).

We note that a biologically feasible range for *R*_0_ and *k* is: 1 ≤ *R*_0_ ≤ 10 and 0 ≤ *k* ≤ 10, thus we use values of these parameters in such range to illustrate our model behavior through simulations. However, the model is general to accommodate any other positive values of such parameters.

Next, we will consider the case if we take *µ* → ∞, which implies *R*_0_ → 1^+^ or *k* → 0^+^, and determine the strain winning in competitive exclusion.

As mentioned, we can rewrite pairwise invasion fitness so that we highlight two opposing terms, where the first one, is completely anti-symmetric in the reverse 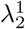, thus contributes only to competitive exclusion.

Whereas, the second term in the square bracket captures the trait variation that may lead to outcomes beyond exclusion. Here, recall that, if we impact on the system so that *R*_0_ → 1 or *k* → 0 to get the phenomenon of exclusion, then 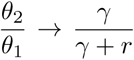, and 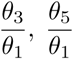 go to 0. Hence, the second part tends to 0 and determination of winner/extinct strain depends on the sign of

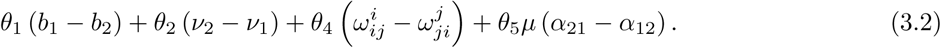

If the sign of this expression is positive then strain 1 will be the winner strain and vice versa. Generally, using these arguments, it is still hard to consider exactly the single winner without computing explicitly the term (3.2). The final answer will depend on the advantage in terms including transmission Δ*b*, duration of carriage Δ*ν*, transmission probability of a strain from a co-colonized host 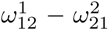 and susceptibility to co-colonization (*α*_21_ − *α*_12_). We will study particular cases in the next sections, but an overview of the range of possible scenarios is given in Table 3.

**Table 3:** Which trait variation between 2 strains leads to which final outcome? Scenarios of variation in biological parameters analyzed with the quasi-neutral coinfection SIS model for 2-strains, and the final ecological outcome of their dynamics. There can be exclusion of strain 1 (E1), exclusion of strain 2 (E2), coexistence (C), and bistability (B). Not all trait variations lead to coexistence. In a majority of scenarios, for a given set of trait variation on ≥ 2 trait axes, the final outcome between two strains may shift with coinfection prevalence in the system.

| Ecology | Exclusion Axis |  |  | Coexistence-Bistability Axis |  | Final outcome for 2 strains |
| --- | --- | --- | --- | --- | --- | --- |
| *Nr. traits varying* | $\beta_i$ | $\gamma_i$ | $p_{ij}^i$ | $\gamma_{ij}$ | $k_{ij}$ | *Possible equilibria with given trait variation* |
| 1 | ✓ | × | × | × | × | E1 or E2 (depends on $R_0^i$) |
| 1 | × | ✓ | × | × | × | E1 or E2 (depends on $R_0^i$) |
| 1 | × | × | ✓ | × | × | E1 or E2 (depends on $\omega_{12}^1 - \omega_{21}^2$) |
| 1 | × | × | × | ✓ | × | E1, E2, C, B (depends on $u_{ij}$) |
| 1 | × | × | × | × | ✓ | E1, E2, C, B (depends on $\alpha_{ij}, (**)$) |
| 2 | • | • | • | × | × | E1 or E2 (*) |
| 3 | ✓ | ✓ | ✓ | × | × | E1 or E2 (**) |
| $\geq 2$ | • | • | • | ✓ | × | E1, E2, C, B (#) |
| $\geq 2$ | • | • | • | × | ✓ | E1, E2, C, B (#) |
| $\geq 2$ | • | • | • | ✓ | ✓ | E1, E2, C, B (#) |

**The role of co-infection (co-colonization) as a potential gradient modulator for system behavior:**
(\*) In this case, for any combination of *fixed* trait variation between two strains (with traits denoted by • varying or not varying), there is only competitive exclusion and up to one shift is possible as a function of $\mu$ ( $I^*/D^*$ ). Thus the system can shift from *Exclusion of 1* → *Exclusion of 2* or vice versa, only once, because of a particular overall coinfection prevalence ( $D/T = 1/(\mu + 1)$ ).
(\*\*) In this case, for any combination of *fixed* trait variation between two strains, only competitive exclusion is possible, but now up to 2 shifts are possible as a function of $\mu$ ( $I^*/D^*$ ). Thus the system can shift three times between opposite exclusion equilibria, because of overall coinfection prevalence.

#### What determines coexistence?

By the previous arguments, in order to have the coexistence of two strains, in our model with coinfection, the essential condition is *R*_0_ *>* 1 and *k >* 0 large enough. Coexistence opportunities can only come from advantages that may arise in coinfection. In other words, this requires the ratio of single to co-colonization *µ* tend to 0. In conclusion, from the analysis until here, we have that:

1. The perturbations only in *β*_*i*_, *γ*_*i*_ and 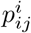 lead to the competitive exclusion. Thus strain-specific transmission and/or clearance rates, and the strain-specific transmission biases from mixed co-colonized hosts only create forces favouring exclusion in the system.
2. The perturbations in co-colonization clearance rates and susceptibilities, *γ*_*ij*_ and *k*_*ij*_, create more complex scenarios including exclusion of each strain, coexistence or bistable exclusion steady states. Thus, only through the possibility of asymmetries in co-colonization (co-infection) parameters can the strains mediate their mutual coexistence.

In the next section, we will consider the phenomena: exclusion, coexistence or bistable exclusion of either strain according to *µ*.

## 4 Effect on each trait variation on final outcome

In this very general version of the model, two strains vary along several fitness dimensions: transmission, clearance rate, co-infection susceptibilities, and possible biases in the clearance of co-infection and transmission from the co-infection compartment. In the following we will explore these dimensions in detail, and what is their effect on the competitive dynamics for *N* = 2.

In the subsections 4.1, 4.2, 4.3, and 4.4, for each case considered, we study the values of mutual invasion fitnesses 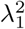 and 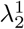 as a function of the ratio of single to co-colonization *µ*. Without loss of generality, in computation we assume that *ν*_1_ < *ν*_2_, hence strain 1 is cleared more slowly than strain 2, giving it an advantage in duration of carriage (Δ*ν >* 0). First, we recall the relative weights (*θ*_*i*_) of each trait in terms of *µ, µ* ∈ [0, +∞) in the Table 1.

### 4.1 Definite drivers of competitive exclusion

In this model, exclusion always results from strain-specific transmission and clearance rate of single infection, and transmission probability from coinfected hosts, if other parameters are equal. Below we explore these three axes of trait variation in more detail. First, we consider variation only in *β*_*i*_, *γ*_*i*_ and 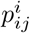. We assume equal parameters for co-colonization clearance *γ*_*ij*_ = *γ* and interaction coefficients between strains *k*_*ij*_ = *k*. The two invasion fitnesses (Figure 2) are:

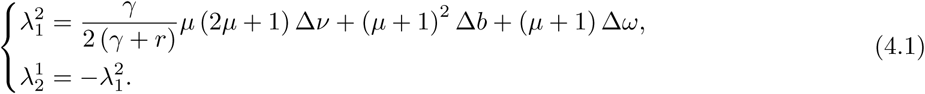

**Figure 2:**
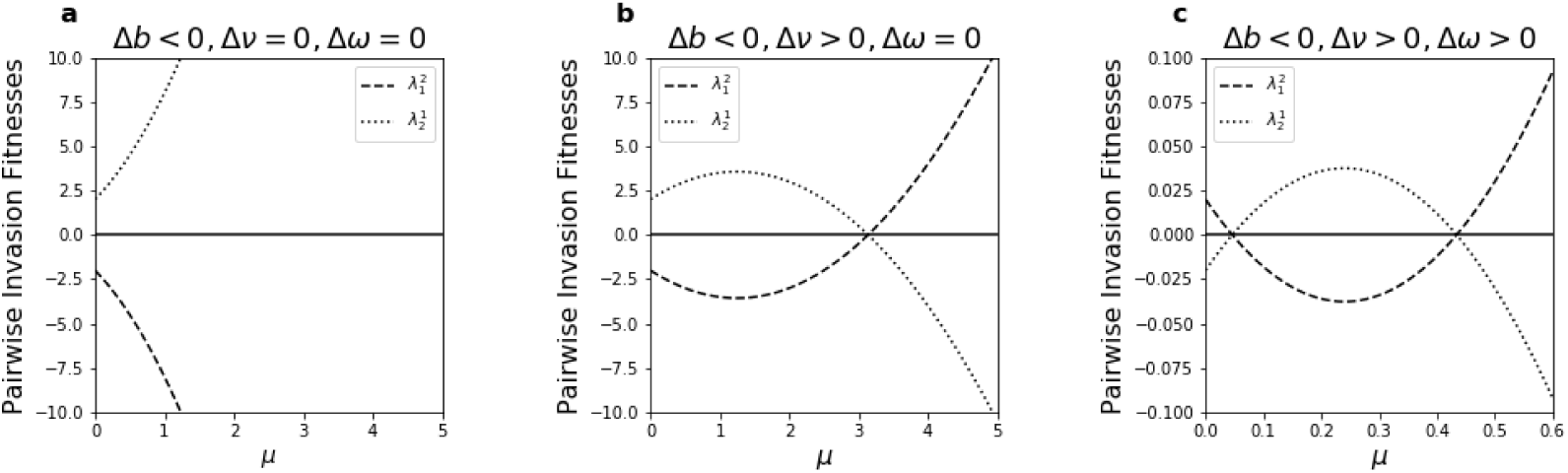
Pure competitive exclusion from variation in three traits, but the winner strain may depend on relative coinfection prevalence (single-to-co-infection ratio *µ* = *I*^∗^*/D*^∗^). Competitive exclusion is the only scenario when two strains vary only in transmission rate *β*_*i*_ and/or infection clearance rate *γ*_*i*_ and/or transmission probability from coinfected hosts 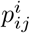 (See (4.1)). Completely anti-symmetric mutual invasion leads to competitive exclusion. The strain with the positive invasion fitness excludes the one with the negative invasion fitness. Here we illustrate the values of 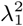 and 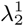 as a function of the ratio of single to co-colonization in the system *µ* = 1*/*(*k*(*R*_0_ *−* 1)), with *γ* = 1.5, *r* = 0.5, in three cases of (**a**) Variations in transmission rates *β*_*i*_ only with Δ*b* = *−*2, (**b**) Variations in transmission rates *β*_*i*_ and clearance rates *γ*_*i*_ with Δ*ν* = 4 and Δ*b* = *−*2, and (**c**) Variations in transmission rates *β*_*i*_, clearance rates *γ*_*i*_ and transmission probability from coinfected hosts 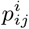 with Δ*ν* = 4, Δ*b* = *−*2 and Δ*ω* = 2.02.

These fitness coefficients are completely anti-symmetric, implying competitive exclusion as the only outcome. Thus it doesn’t matter that there is co-infection in the system (*k >* 0). For promoting coexistence, this is not sufficient by itself. Variable co-infection susceptibilities or traits between strains would be an additional requirement. When strains behave equally in all processes related to coinfection, coinfection cannot rescue them from the destiny of competitive exclusion. However, as we explore below, overall prevalence of coinfection can actually shift between the winning and losing strain. This can happen only if there is variation in duration of carriage. For example noticing that when *β*_*i*_ is the only trait varying between two strains, *k* does not appear in *θ*_1_, this indicates that transmissibility’s variation, uniquely determines the winner between two strains; its relative contribution to 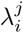 cannot be altered by coinfection.

If variation is only in transmission rates *β*_*i*_, then the strain with bigger *β*_*i*_ excludes the other one, see figure 2 **(a)**. This fact holds for *N* strains in general and is proved in (Le et al., 2021).

If a strain is superior in both fitness dimensions, which means it has smaller clearance rate and greater transmission rate (Δ*b >* 0, Δ*ν >* 0), then for all value of *µ* or *R*_0_, it surely will be the winner par excellence, which can be easily seen from (4.1). This is unsurprising and naturally expected.

However, if a strain is better in one trait but worse in another, for example if the strain with longer duration of carriage (lower clearance) also has smaller transmission rate, the determination of the winning strain depends on value of *µ* (and in general also *R*_0_), see Figure 2b.

Since we have fixed here *γ* = 1.5, *r* = 0.5 by convention, if we fix neutral transmission rate *β*, the value of *µ* now depends only on the mean interaction coefficient in co-colonization *k*. This means, when *k* is high, thus when strains tend to allow each-other more in co-colonization, *µ* is sufficiently close to 0, strain 2, which has the smaller transmission rate but longer duration of carriage is the winner. In contrast, when *µ* is larger, thus when hosts are less vulnerable to co-colonization, the strain 1, which has bigger transmission rate and smaller duration of carriage is the winner.

This illustrates how the relative advantage between two strains, differing in two traits, depends on coinfection prevalence.

It is interesting to note that, if variations are only in two of which including transmission rates *β*_*i*_, clearance rates *γ*_*i*_ and transmission probability from coinfected hosts 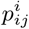, we can have at most one shifting outcome, i.e. shifting once from the exclusion of one strain to the exclusion of the other one, see proof in S3.1.

Following the same analysis, we can study the model in which transmission probability from co-colonized hosts, denoting within-host advantage, 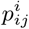 displays strain-specific perturbation. We note that if there is variation in 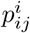 only, if 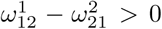, strain 1 excludes strain 2, and vice versa. This can be understood via the precedence advantage that one strain has from mixed coinfected hosts if it arrives first, and thus gets transmitted more.

If there are combinations of variations in within-host transmission advantage 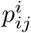, as well as other traits *β*_*i*_ and/or *γ*_*i*_, the final outcome is more complex (see Table 3 and Figure 2 **(c)**). However, the long time competitive result is always exclusion of one strain from the system. If perturbations occur in transmission rate *β*_*i*_ and transmission probabilities from mixed coinfection 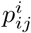, the winning strain depends on coinfection prevalence in the system, (*µ* = *I*^∗^*/D*^∗^) because 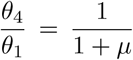. If perturbations occur in duration of single carriage (i.e. strain-specific clearance rates *γ*_*i*_) and 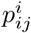, the winning strain depends on more strain-transcending parameters: *µ, γ* and *r*. This can be explicitly observed in the ratio 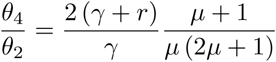, which ultimately affects the signs of pairwise invasion fitnesses. Figure 2 **(c)** shows us a special example in which the exclusion of either strains shifts twice when *µ* varies from 0 to ∞.

### 4.2 Four scenarios possible with variable co-infection clearance rates *γ*_*ij*_

As we can see from the fully explicit expression of pairwise invasion fitness, in this model, the coinfection clearance rate axis contributes to 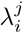 with a term 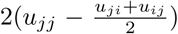. Thus what matters is the comparison between clearance rate of same strain coinfection vs. the mean coinfection clearance rate of mixed-strain coinfection. While there are no restrictions for how these can vary, in the following we consider three special cases where the variation in coinfection duration depends on variation in single infection duration. We also assume no variation in transmission probability from coinfected hosts 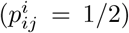 but the results for the general case can be easily derived using Eqs.(3.1).

#### 4.2.1 Case 1: Unbiased clearance in mixed carriage 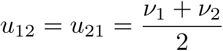

In this first case, we consider that the clearance rate of mixed carriage is unbiased and equal to the mean of the two clearance rates of single colonization. We have *u*_11_ = *ν*_1_, *u*_22_ = *ν*_2_ and the invasion fitnesses between two strains are anti-symmetric:

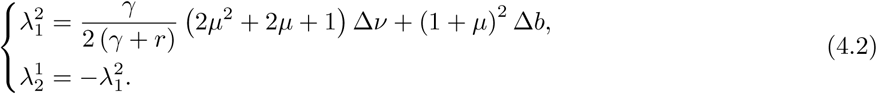

This case leads again to the pure competitive exclusion. Whichever strain has positive 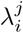 will be the winner.

Similar to the case in section 3.1, if a strain is superior in both transmission *β*_*i*_ and clearance *ν*_*i*_, it will be the winner.

However, if Δ*ν* and Δ*b* have opposite sign, meaning one strain has advantage in clearance and the other has advantage in transmission, the final winner will depend on coinfection prevalence, hence on *µ*. From the formula of the invasion fitness, it can be seen that in this system, the clearance rate differential (i.e. in duration of carriage) has more important role than the transmission rate difference in helping an inferior strain overcome and overturn its fitness disadvantage as *µ* increases.

#### 4.2.2 Case 2: Decreased clearance in mixed carriage *u*_12_ = *u*_21_ = min{*ν*_1_, *ν*_2_} = *ν*_1_

Here, we explore the case when mixed co-infection clearance rate corresponds to the minimum of the two single infection clearance rates. Without loss of generality, we assume that Δ*ν* ≥ 0 i.e. *ν*_1_ ≤ *ν*_2_. We still have *u*_11_ = *ν*_1_, *u*_22_ = *ν*_2_, but in hosts carrying a mixture of strain 1 and strain 2, this creates an advantage in co-infection for the opponent strain (the one with the faster strain-specific clearance). The two invasion fitness coefficients are not anti-symmetric anymore, hence allowing for more scenarios beyond exclusion (see figures 3 **(a**,**b)**):

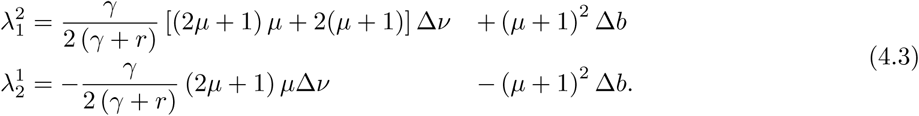

**Figure 3:**
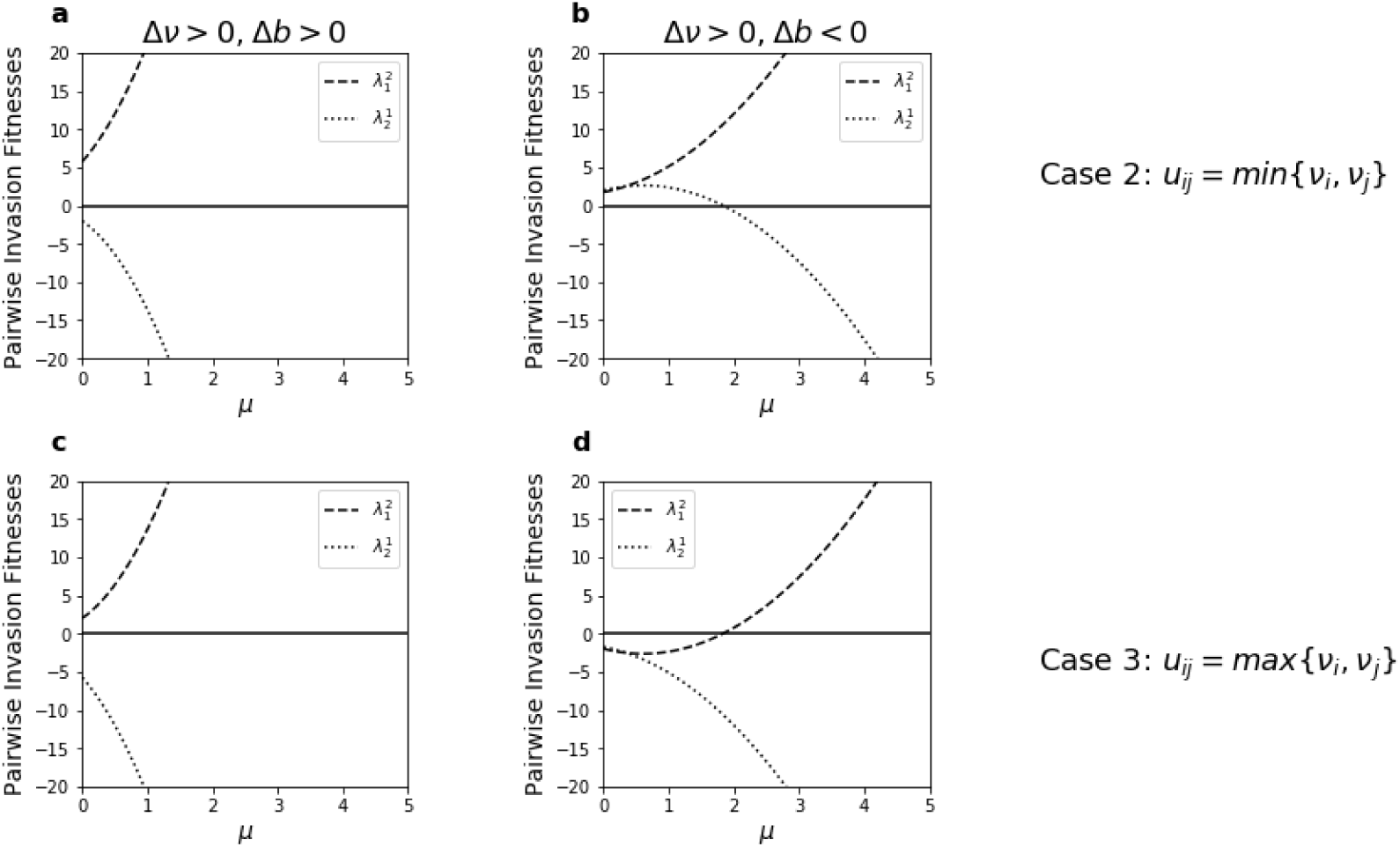
Competitive exclusion can be broken with variable coinfection clearance rates, and the result may depend on *µ*. We illustrate possible scenarios resulting from deviation from symmetry in the mixed coinfection clearance rate *γ*_*ij*_ as a function of the ratio of single to coinfection *µ*. **a-b** Coinfection clearance equals the minimum clearance rate of either strain: *u*_12_ = *u*_21_ = min*{ν*_1_, *ν*_2_*}* = *ν*_1_ (see (4.3)). **c-d** Coinfection clearance equals the maximum clearance rate of either strain: *u*_12_ = *u*_21_ = max*{ν*_1_, *ν*_2_*}* = *ν*_2_ (see (4.4)). We choose *γ* = 1.5 and *r* = 0.5 as in figure 2. For each sub case, we plot the mutual invasion fitnesses for transmission advantage and disadvantage of strain 1, respectively: Δ*b >* 0 (**a**,**c**) and Δ*b* < 0 (**b**,**d**). In particular, in the first column Δ*b* = 2, and in the second column Δ*b* = *−*2. The clearance rate differential Δ*ν* is assumed Δ*ν* = 5 attributing higher duration of carriage to strain 1. In the first column, strain 1 is superior in all fitness dimensions, and coinfection clearance cannot overturn the result. In the second column, strain 1 is not superior in all fitness dimensions, and coinfection matters for the final result.

If one strain, denoted to be strain 1 in Figure 3a, has larger transmission rate and lower clearance rate (case when Δ*b >* 0, Δ*ν >* 0), it will again be the only survivor for all *µ*.

However if the advantage is only in one of the two traits (case when Δ*b*, Δ*ν* have opposite signs), for example the strain with smaller clearance rate has lower transmission rate, as in Figure 3b, then coexistence can occur. However, even in this situation, coexistence can only be possible for sufficiently low *µ*, i.e. *k* large enough for fixed *R*_0_ (we already fix *γ* and *r*). This means that increasing the relative prevalence of co-colonization or coinfection in the system, via higher facilitative interactions, can promote coexistence of two strains. This links back to the arguments at the end of section 2.

The phenomenon of coexistence arising here is similar to what has been found before in the context of virulence evolution (Alizon, 2008), where decreased clearance in coinfection was observed to promote coexistence, hence persistence of more virulent strains (here strain 2 if Δ*ν >* 0, and in the limit of strain 2-*R*_0_ below 1).

#### 4.2.3 Case 3: Increased clearance in mixed co-infection *u*_12_ = *u*_21_ = max {*ν*_1_, *ν*_2_} = *ν*_2_

The co-infection clearance rate in mixed carriage here is assumed to be equal to the maximum value of the strain-specific clearance rates in single infection. As in the previous section, we still assume that Δ*ν* ≥ 0 i.e. *ν*_1_ ≤ *ν*_2_. We have *u*_11_ = *ν*_1_, *u*_22_ = *ν*_2_ and the pairwise invasion fitness coefficients are (see figures 3 **(c**,**d)**):

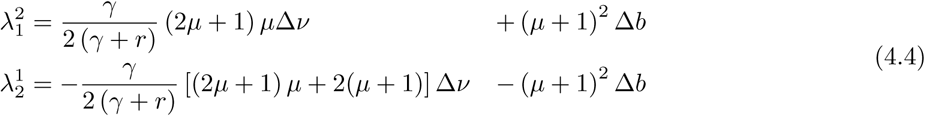

If strain 1 has smaller clearance rate and larger transmission rate, it is again the superior strain in the system, independently of coinfection parameters, like previous cases (Figure 3c).

However, if strain 1 has smaller clearance rate but also lower transmission rate, bistability of exclusion can occur when *µ* is small enough. When *µ* becomes larger and tends to infinity, we obtain only competitive exclusion, as mentioned earlier and proven in Section 2. In that extreme, strain 1 is the only persistent strain over long time (Figure 3d).

In conclusion, by the explicit formulae of 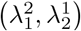 in the cases above, we can also prove that for *µ* large enough, the strain which has smaller clearance rate will be the only strain persisting in the system.

### 4.3 Four scenarios from variation in pairwise co-colonization susceptibilities *k*_*ij*_

Another fitness dimension is how the strains facilitate or compete in altered susceptibilities to co-infection via the coefficients *k*_*ij*_. Above we assumed they are all equal to the reference *k*. But when variation in this parameter is allowed, as shown already in (Gjini and Madec, 2017), all four ecological scenarios are possible, and thus the effect is to open up space for coexistence and bistability among two strains, when competitive exclusion is expected from other parameters. According to the derivation of the reduced model in (Madec and Gjini, 2020), the perturbations in co-colonization interaction matrix for *N* = 2 satisfy 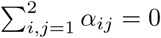 when *k* is defined by the mean of *k*_*ij*_. However, without loss of generality, one can shift the *α*_*ij*_ by the same constant, without changing the mutual 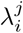 and consequently without changing the dynamics. The explicit formulas for two pairwise invasion fitnesses are (see figure 4)

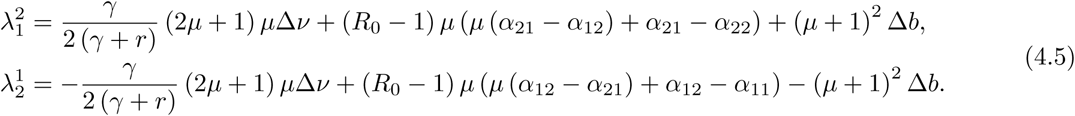

**Figure 4:**
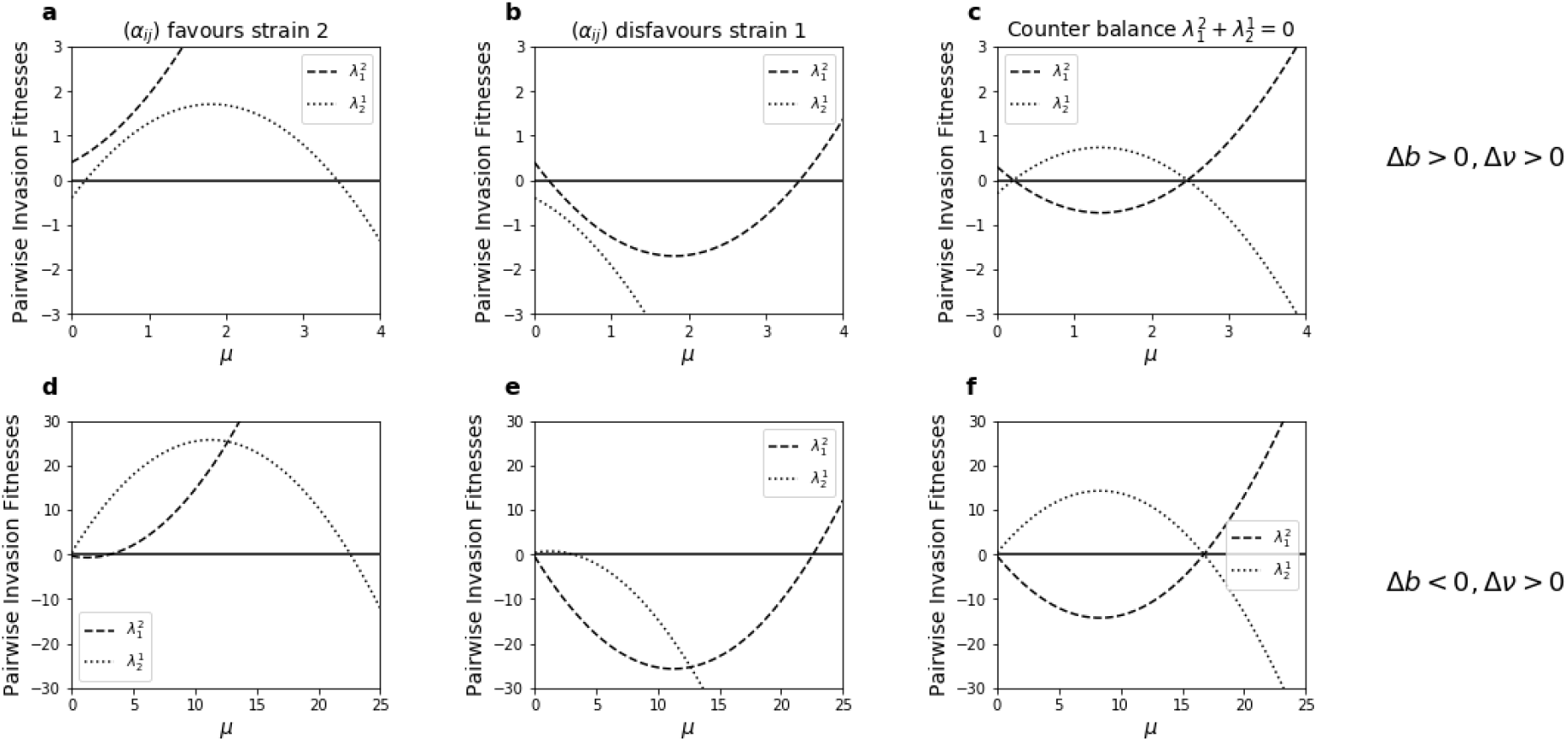
Breaking the competitive exclusion with co-colonization interactions *k*_*ij*_ (see Eqs (4.5)). We compute pairwise invasion fitnesses 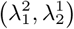 according to *µ* in various cases of co-colonization interaction matrix (*α*_*ij*_) with *R*_0_ = 5, *r* = 0.5 and *γ* = 1.5. **(a-c)** We illustrate the cases of transmission superiority of strain 1: Δ*b >* 0, when Δ*b* = 0.4, Δ*ν* = 0.8. In **(d-f)** we plot 2-strain invasion fitnesses for transmission superiority of strain 2: Δ*b* < 0, when Δ*b* = *−*0.4, Δ*ν* = 0.8, with the same *γ, r* and *R*_0_ as in **(a, b, c)**. Coinfection clearance rate *γ*_*ij*_ is assumed equal to *γ* and transmission probability from coinfected hosts carrying a mixture of two strains 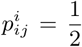. Subplots with the same values (*α*_*ij*_) lie in the same column. In particular, we consider 3 structures: **(a, d)** 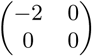 (Eqs (4.6)); **(b, e)** 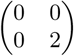 (Eqs (4.7)); **(c, f)** 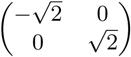 (Eqs (4.8)) for variation in co-colonization interactions. Except for when the *α*_*ij*_ exactly counterbalance effects on 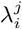 (c,f) there is potential for more scenarios beyond competitive exclusion, induced by coinfection susceptibilities between strains.

In this spirit, below we consider a few special cases of *α*_*ij*_ variation between strains, to highlight the effect of co-infection susceptibilities when they provide:

1. **an advantage to strain 2:** 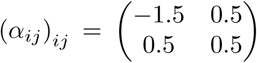, whose effect on 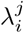 is equivalent to 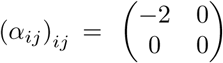, and increases relatively 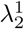:

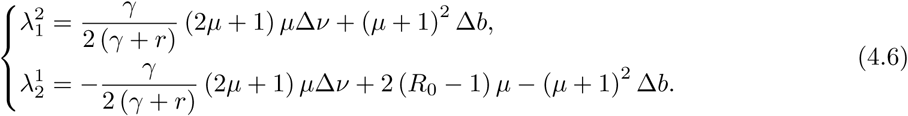
2. **a disadvantage to strain 1:** 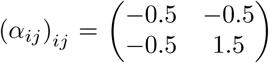; equivalent to 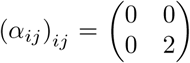 when shifted by the appropriate constant, which relatively decreases 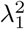:

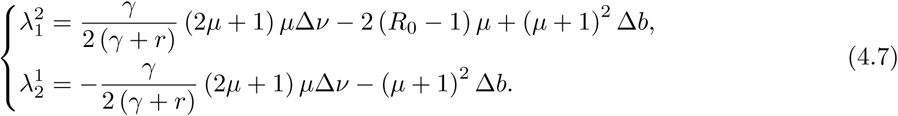
3. **and exactly counterbalanced effects on either strain:** 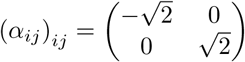, whose impact on 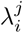 is to decrease 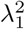 and increase 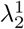 by exactly the same amount:

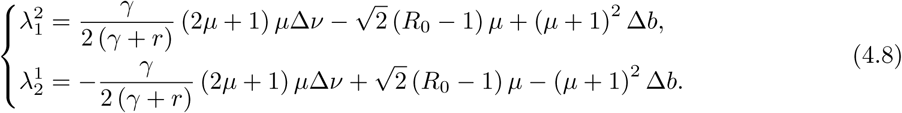

In Figure 4, we consider these cases, where besides *k*_*ij*_, we allow also variation in transmission *β* and clearance *γ* of each strain, but assume initially symmetry in other traits. We assume Δ*ν >* 0 (strain 1 is cleared more slowly). In Fig. 4a-c we illustrate competitive outcomes (dependent on mutual signs of 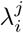), when strain 1 is superior in transmissibility, and in Fig. 4d-f we show outcomes when strain 2 is superior in transmissibility instead. Naturally as *µ* → ∞ the role of coinfection interaction asymmetries vanishes, and the system tends to exclusion, but for low values of *µ*, the structure of the *α*_*ij*_ matters. In particular, if it is asymmetric (Figure 4,a-b, d-e) there can be at most 3 scenarios as a function of *µ*: exclusion - coexistence -exclusion, or exclusion - bistability-exclusion. In particular increase in 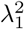 acts to enable coexistence when Δ*b >* 0, and a decrease in 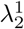 acts to enable bistability when Δ*b* < 0. Whereas, if co-colonization interactions have exactly counterbalanced effects on 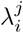 (Fig. 4c,f), there can only be alternating exclusion scenarios as a function of *µ*.

### 4.4 Adding variation in transmission probability from coinfected hosts

In this case, besides transmission and clearance rates variations (Δ*b*, Δ*ν*), and co-colonization susceptibilities (*α*_*ij*_), we add to the system variation in terms of a slight priority effect between strains for transmission from coinfection Δ*ω* ≠ 0. The explicit formulae for pairwise invasion fitnesses of two strains are as follows:

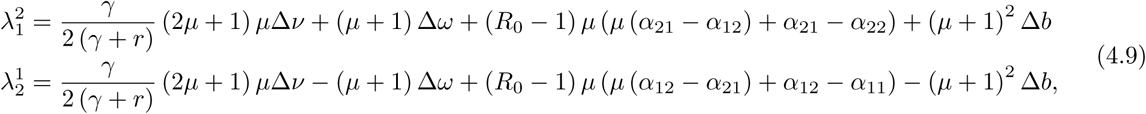

Recall that 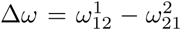 represents the relative advantage of strain 1 from arriving first within-host, in transmission from mixed coinfection. It is clear from the expression above, that when variation in this within-host advantage is combined with variation in coinfection clearance rates or co-colonization susceptibility factors between strains, coexistence and bistability also become possible. In this case within-host and between-host competition combine to give rise to different outcomes. In figure S6 we illustrate the effect of Δ*ω* on the baseline outcomes of Figure 4a-c. The effect of Δ*w* < 0 is to increase the potential for strain 2-only competitive exclusion and coexistence with strain 1 in the system, oftentimes overturning the baseline result especially so if *µ* small. The importance of transmission biases from mixed coinfection Δ*w* is unsurprisingly higher when relative coinfection prevalence is higher in the system.

### 4.5 The qualitative outcome for two strains can shift multiple times with *µ*

Until now we have seen the important and explicit role of strain-transcending parameters, (e.g. *R*_0_, *k*, and specifically *µ*) which define the core neutral system of this model, on the ultimate competitive outcome between strains at the epidemiological level. We have seen that the same relative variation between strains, displayed in (Δ*b*, Δ*ν*,…) will have a different impact in a system with larger or lower overall prevalence of coinfection, relationships that are completely transparent in the 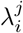. Sometimes the effect will be quantitative, changing only the speed of dynamics without affecting the 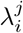 signs (see Supplementary figure S2). Other times the effect will be qualitative, changing the signs of the pairwise invasion fitnesses and hence the dynamics. Furthermore, using such full analytic transparency, we can also prove mathematically special results to make the claims about qualitative shifts more precise. Depending on how many traits and which traits vary in the system, we can have at most one, two, or more shifts with *µ* (see Table 2). For example we can prove that with variation only in transmission and clearance rates, there can be at most one shift in final outcome as a function of *µ* (Text S3.1)

A very special case arises, when the same system can shift 4 times as a function of *µ*. We have proven that a necessary condition for its occurrence is the presence of variation in both coinfection clearance rates *γ*_*ij*_ and vulnerabilities to coinfection *k*_*ij*_. If any of these is missing, 4 shifts as a function of *µ* are impossible (see Text S3.2 for the formal proof). An illustration of such a special case is given in Figure 5, where the system traces all 4 quadrants as *µ* goes from 0 to ∞, highlighting an extreme case of the critical role of coinfection, for the relative hierarchical advantages between two strains and their selection dynamics. We see that the system is characterized by competitive exclusion of strain 2 for *µ* low, then tends to coexistence of both strains for increasing *µ*, followed by competitive exclusion of strain 1 for even higher *µ*, until it returns again in the competitive exclusion region of strain 2 as *µ* → ∞.

**Figure 5:**
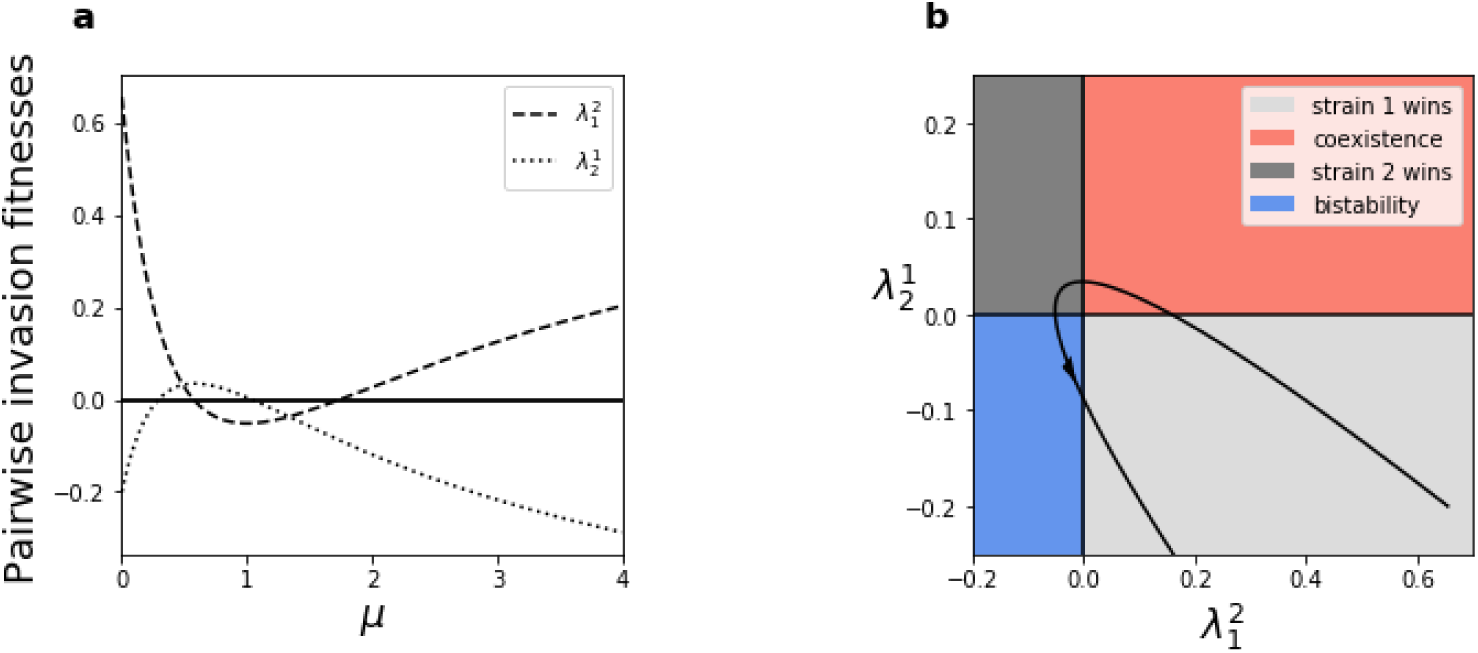
Four ecological scenarios may happen depending on *µ* (single-to-co-infection ratio) under fixed trait variation. Variations here are in transmission rates, infection and coinfection clearance rates, and co-colonization susceptibilities between strains. The strain-transcending parameters are assumed *γ* = 2, *r* = 0.2 and *R*_0_ = 2.5. We assume that Δ*b* = 0.2, Δ*ν* = 0.5, the coinfection clearance rate *γ*_*ij*_ with *u*_*ij*_ = min*{ν*_*i*_, *ν*_*j*_ *}* and values of (*α*_*ij*_) to be 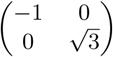. The ultimate ecological outcomes when *µ* goes from 0 to *∞* include: exclusion of strain 2, coexistence, exclusion of strain 1, bistablity, then back to the exclusion of strain 2. Figure **(a)** plots two pairwise invasion fitnesses as function of *µ*. Figure **(b)** plots the parametric curve 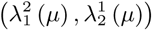 for *µ* in the same range in figure **(a)**, from 0 to 4. This cure crosses all four quadrants, which means we have all four outcomes when *µ* varies.

### 4.6 Parameter regions for 4 outcomes in our SIS model with co-infection

Until now, we have considered fixed trait variations between two strains, and varied *µ* to show how their net competitive dynamics driven by 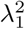 and 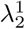 will depend on the ratio of single to coinfection in the neutral system. This context is defined by basic reproduction number *R*_0_, and *k*, of the neutral model (see Table 1). Next, we consider the distribution of 4 possible ecological outcomes across different systems, as a function of *R*_0_ and *k*. In figure 6 we represent the long time behaviour of a 2-strain model with perturbations in transmission rate *β*_*i*_, clearance rate of single colonization *γ*_*i*_ and clearance rate of co-colonization *γ*_*ij*_ with *u*_11_ = *u*_12_ = *u*_21_ = *ν*_1_ and *u*_22_ = *ν*_2_. Figure 6 shows which combinations of Δ*b* and Δ*ν*, lead to one of the four scenarios: exclusion of strain 1 or 2, coexistence or bistable state, for each *R*_0_ and *k*, for assumed symmetry in *k*_*ij*_ and in 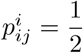.

**Figure 6:**
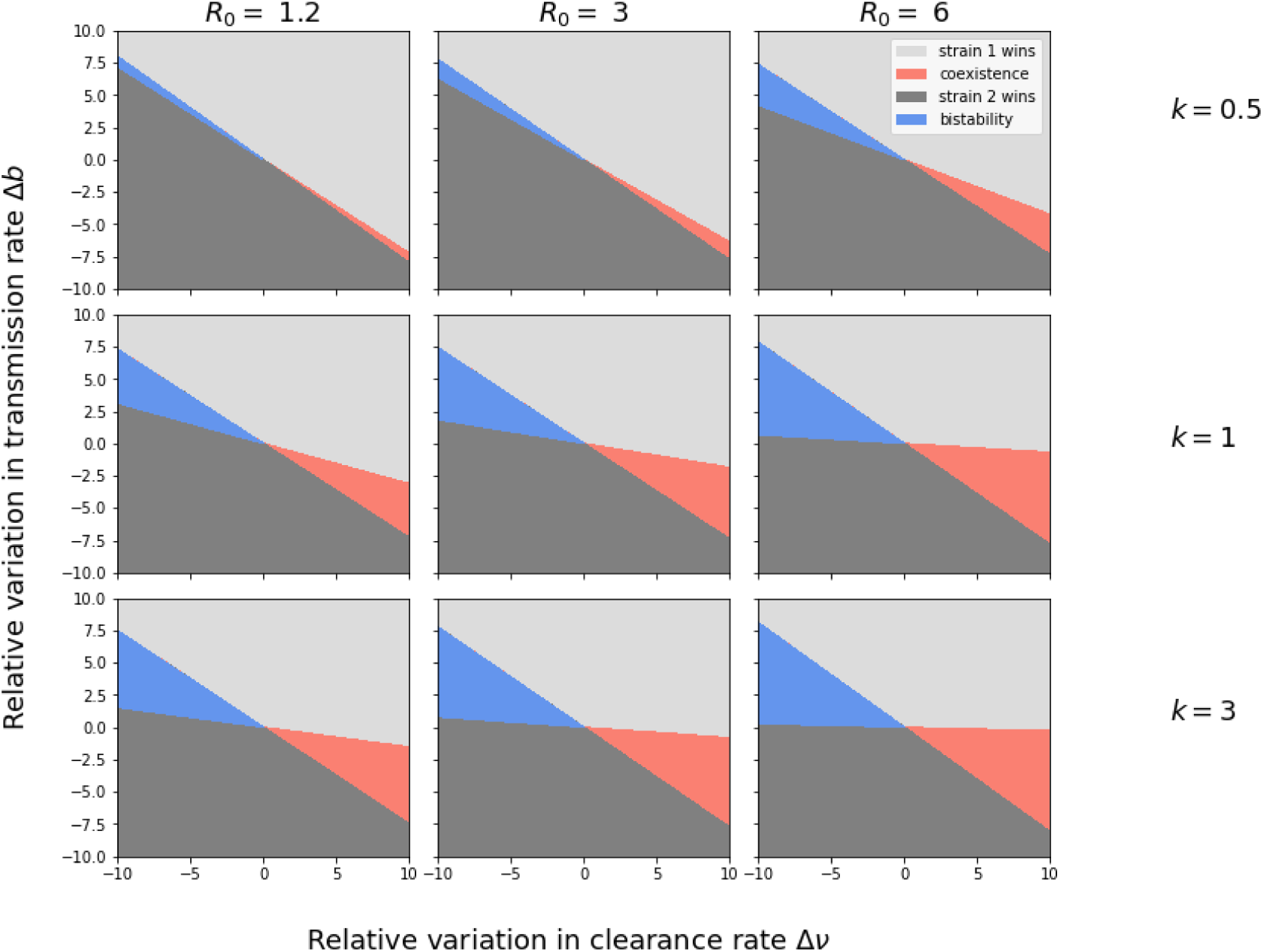
Illustration of 4 possible outcomes, as a function of relative variation in transmission and clearance rate between two strains, for different values of *k* and *R*_0_. We highlight the respective regions in different colors, according to the critical relationship between Δ*b*, Δ*ν, k* and *R*_0_ when perturbations happen only *β*_*i*_, *γ*_*i*_, and mixed coinfection clearance happens base on strain-specific clearance rates *γ*_*ij*_ with *u*_11_ = *u*_12_ = *u*_21_ = *ν*_1_ and *u*_22_ = *ν*_2_. We choose the values *γ* = 1 and *r* = 0.2. Recall that the higher Δ*ν >* 0, the higher the advantage of strain 1 in the system; and the higher Δ*b >* 0, the higher the advantage of strain 1 in the system. We observe coexistence and bistability arise only when the disadvantage in one trait is compensated by an advantage in the other. In particular coexistence is enabled when the disadvantaged strain 2 in clearance rate benefits from reduced clearance in mixed co-colonization with strain 1.

It can be seen that we can choose suitable values of relative trait differences Δ*b* and Δ*ν* to observe a given scenario, typically coexistence and bistability arise for relative advantage in one trait and relative disadvantage in the other, hence a trade-off between transmission and clearance. Notice that when *R*_0_ or *k* become larger, increasing relative coinfection prevalence in the corresponding neutral system, (i.e. reducing *µ*), the possibility for coexistence or bistability expands in the system. Near the origin, it’s harder to obtain coexistence or bistable state rather than competitive exclusion.

The results of Figure 6 can be used to connect our system’s behavior to strain-specific basic reproduction numbers 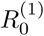 and 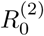. In a model with coinfection, strain-specific *R*_0_ are not sufficient to determine the long-time epidemiological competition between two strains. We show in Text S1 and in Figure S1 that even for the same values of *R*_0,1_ and *R*_0,2_ we can have different long-time scenarios between the two strains. The result in coinfection depends on the particular combination of traits, strain-specific transmission and clearance rates. In particular, even for the same strain-specific *R*_0_’s, a system with bigger variation in clearance rate between two strains is more sensitive to coinfection, and it is where coinfection parameters can shift the dynamics more easily away from competitive exclusion (Fig. S1a). This is because of the advantage conferred to both strains by staying longer in mixed compartment *I*_12_, where from they can have equal chance of transmission.

In Fig. 6 coinfection susceptibilities were assumed symmetric and coinfection clearance rate for mixed carriage was assumed biased toward the same clearance rate of single infection by strain 1: *u*_11_ = *u*_12_ = *u*_21_ = *ν*_1_ and *u*_22_ = *ν*_2_. In another case (Figs. S3-S4), when we remove the bias in coinfection clearance assuming *u*_12_ = *u*_21_ = 0, but allow variation in susceptibilities to coinfection, we see only three scenarios emerge, over all Δ*b* and Δ*ν*. In this case, depending on the structure of *α*_*ij*_, we observe either coexistence or bistability as a third possibility flanked by the two opposite exclusion steady states.

## 5 How trait mean and variation impact coexistence frequencies

Next, we zoom in from criteria for coexistence to the details of the coexistence equilibrium in a system with two strains that vary along multiple fitness dimensions. The general formula in terms of pairwise invasion fitnesses in Eqs. (3.1), when this equilibrium exists, is given by

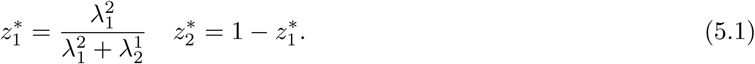

This allows explicit computation for any combination of strain-specific and strain-transcending parameters in the system. Below we focus on the case of variation in transmission rate *β*_*i*_, clearance rate of infection *γ*_*i*_ and in interaction coefficients via susceptibilities to coinfection *k*_*ij*_. We assume the special case of *u*_*ij*_ = 0 and initially no transmission biases in coinfection 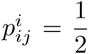 (although we relax this later). We consider only coexistence regimes (Figure 7) and illustrate the equilibrium frequency of one of the strains (here 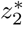), as a function of trait mean and variation between strains. Using the explicit formula 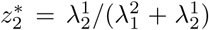, we explore this quantity numerically for two values of mean co-infection susceptibility: *k* = 1.5 and *k* = 0.2, and symmetric cross-strain interactions *α*_12_ = *α*_21_ *> α*_11_ = *α*_22_. We consider 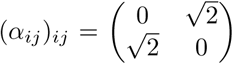 but the general formula for other cases is straightforward (see Text S3). The criteria enabling coexistence are derived below.

**Figure 7:**
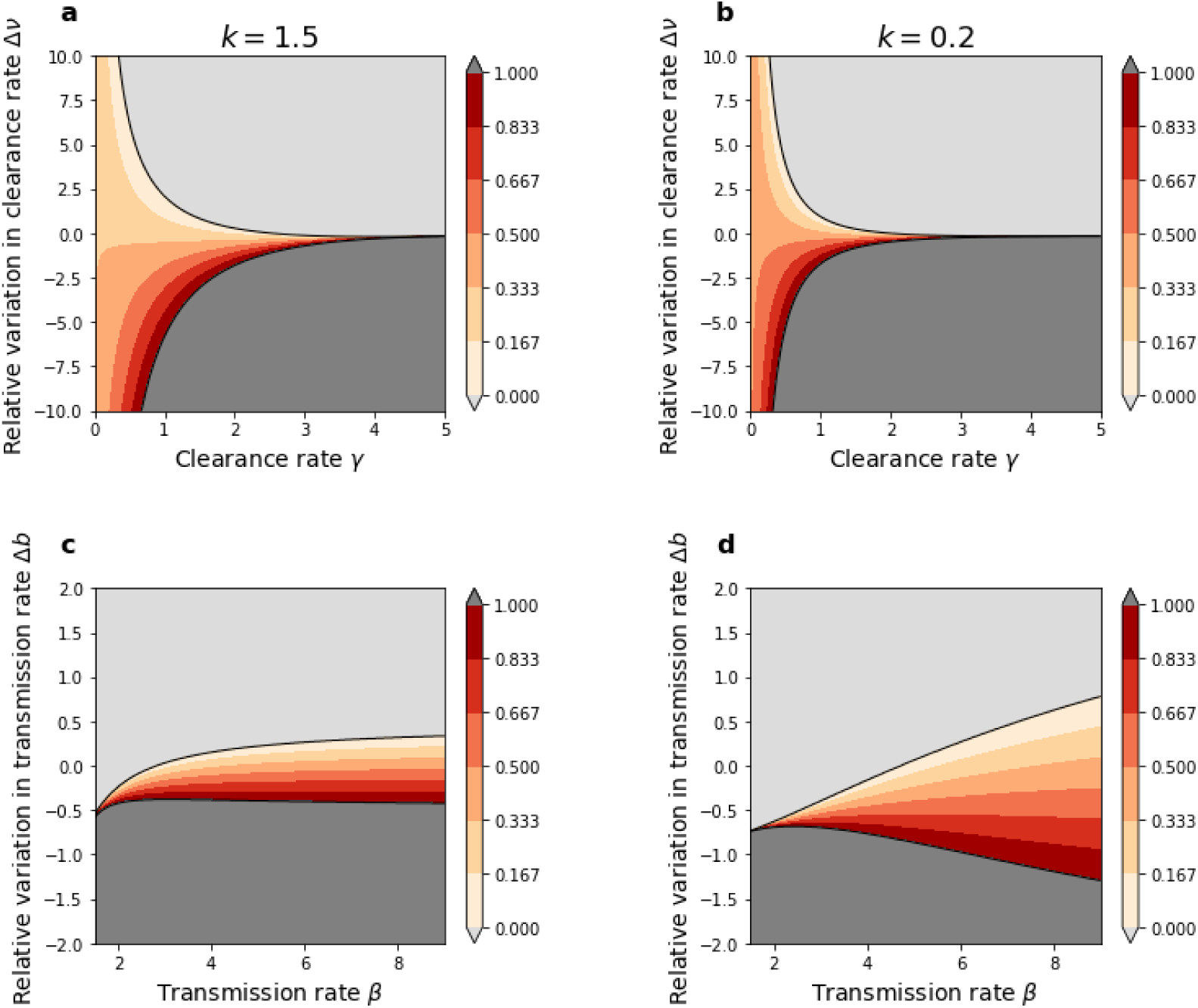
Strain coexistence frequencies depend on a critical interplay between trait mean and variation. We plot the coexistence equilibrium frequency of strain 2 (intermediate shading), as a function of (*β*, Δ*b*) and (*γ*, Δ*ν*) for higher coinfection prevalence (left) and lower coinfection prevalence (right). **a-b. Effect of clearance rate**. In (a-b) we assume *β* = 5.3 and Δ*b* = 0.15. In these figures, we vary *γ* in 0 *≤ γ ≤* 5 and *β* = 5.3, to make *R*_0_ *>* 1. **c-d. Effect of transmission rate**. In (c-d) we assume *γ* = 1 and Δ*ν* = 1 as fixed. We vary *β* between 1.5 and 9, ensuring *R*_0_ *>* 1. Right-column subplots reflect a system with more strain competition preventing co-infection, hence lower coinfection prevalence (*µ* higher) than the left-column subplots. The light grey region represents the exclusion of strain 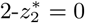 and the dark grey region is for exclusion of strain 1 from the system i.e. 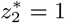. We choose *r* = 0.3 and the matrix (*α*_*ij*_) to be 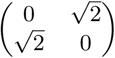 thus unbiased in terms of favouring either strain. The co-infection clearance rate is also assumed unbiased, and equal to the mean. The black lines denote the border lines for which coexistence is no longer possible and the system shifts to either exclusion of strain 1 (yellow) or exclusion of strain 2 (blue). In a-b, the lines are denoted by *T*_1_ (*γ*) and *T*_2_ (*γ*), and given by (5.3) and (5.4). In c-d, the graphs of *S*_1_ (*β*) and *S*_2_ (*β*) are hyperbolic, given by explicit equations (5.6) and (5.7).

### 5.1 Mean and variation in clearance rate of single infection

By (4.3), the equation of boundary 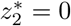 is equivalent to 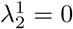 which can be written explicitly as:

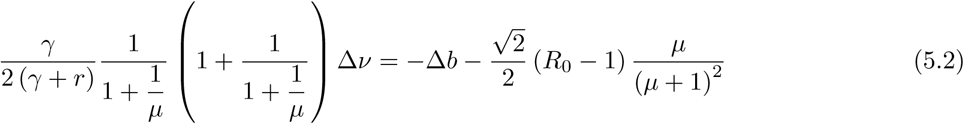

which, by substituting 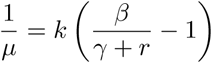, becomes:

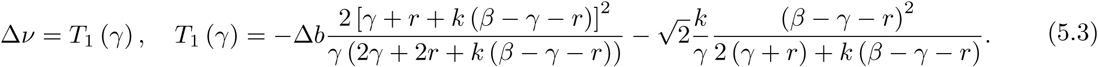

Analogously, we can compute the equation of boundary 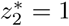, thus 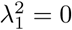, which is

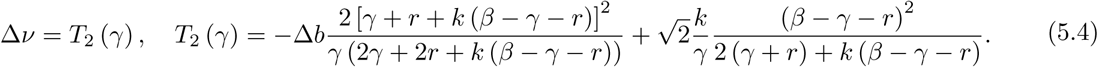

Combining (5.3) and (5.4) we have the condition for 2-strain coexistence between these boundaries, expressed as an inequality for the variation Δ*ν* dependent on the mean *γ*:

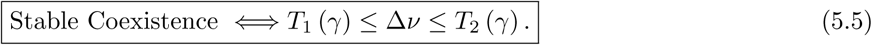

The figures 7a and 7b show critical interplay between mean and variation clearance rates of infection *γ*_*i*_. The light color region in this figure is the region displays coexistence phenomenon, corresponding to the space between two curves which present equations (5.3) and (5.4). The contour shading corresponds to the relative frequency of strain 2. In these figures, we can observe that by decreasing mean co-infection efficiency *k*, we can decrease the possibility of coexistence at the same variation in clearance rate Δ*ν* (7b compared to 7a). For *γ* small enough, which increases overall prevalence in the system, less values of Δ*ν* lead to coexistence. When Δ*ν* becomes larger, the relative frequency of strain 2, decreases.

### 5.2 Mean and variation in transmission rate

Similarly to (5.3) and (5.4) for mean and variation in clearance rate (duration of carriage), and the same assumptions about the other parameters, we can find the equation of boundary 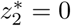 as a function of (*β*, △*b*) as follows:

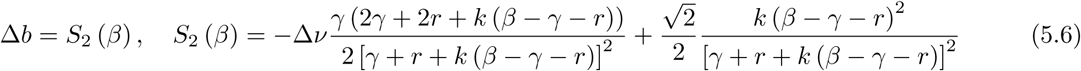

The equation of the boundary 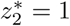 as a function of (*β*, △*b*) is:

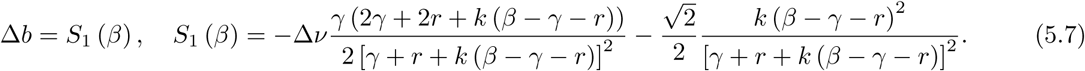

Combining (5.6) and (5.7), we have the condition on intermediate values of Δ*b* that allow 2-strain coexistence:

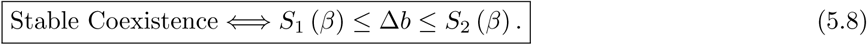

Two figures 7c and 7d show the value of strain 2 in case of coexistence as a function of mean and variation in transmission rates Δ*b*. In contrast to 7a and 7b, by decreasing co-infection vulnerability *k*, we increase the possibility of coexistence at the same variation in transmissibility between strains Δ*b*. This also means that for larger *β*, more values of Δ*b* lead to coexistence. When Δ*b* becomes larger, increasing the transmission advantage of strain 1, the equilibrium frequency of strain 2, decreases. This is similar to Figure 7a and 7b.

### 5.3 Coexistence strain frequencies are explicit

While the entire frequency dynamics for each strain and for all time (*z*_*i*_(*τ*)) is fully explicit in the system, via the replicator equation, so is also the final equilibrium. The value of the equilibrium frequency of strain 2, 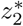 under the coexistence regimes studied above is given by:

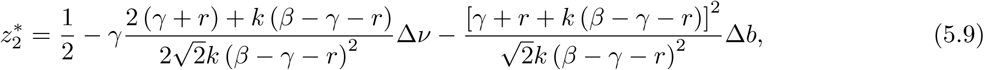

and can be seen to be an explicit function of mean parameters (e.g. *β, γ*) as well as variation between strains (Δ*b*, Δ*ν*). Here, because coinfection interactions are assumed symmetric, it becomes obvious that any differences between strains will make the frequency deviate from the expected frequency of 1/2 under balancing selection. If Δ*ν*, Δ*b >* 0, then strain 1 has an absolute advantage and *z*_2_ will always be inferior than 1/2, if it coexists. However, if the advantage is only in one trait and not in the other (Δ*ν*, Δ*b* of different signs), then strain 2 can increase its equilibrium frequency above 1/2.

More generally, for the rescaled co-colonization susceptibilities matrix being symmetric and satisfying: *α*_*ii*_ = *α*_*jj*_ = *α*_11_ and *α*_*ij*_ = *α*_*ji*_ = *α*_12_, we can write the expression for strain 2 equilibrium frequency, more compactly as a function of *µ, k* and *R*_0_ constituent parameters, mean transmission rate *β*, clearance rate *γ* and average host lifespan 1*/r*:

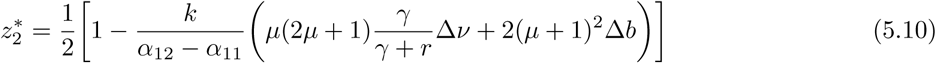

One can notice that variation in each trait Δ*b* and Δ*ν* have their own distinct nonlinear scaling factors for how they impact on ultimate strain success at the epidemiological level, depending directly on the prevalence of co-colonization in the system via the parameter *µ* = *I*^∗^*/D*^∗^. We can immediately see from this formula, how the predicted coexistence level among two strains is attributable to clear mechanisms and clearly identifiable biological differences between strains, which are explicitly weighted by epidemiologic constants.

Even more generally, if in addition there are biases in strain transmission probabilities from co-colonized hosts carrying a mixture of strains, recalling that 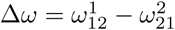 the formula (5.10) now reads

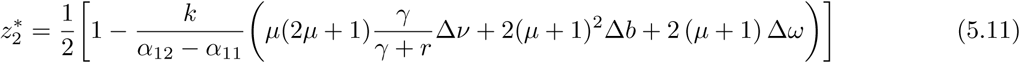

where we can straightforwardly see the contribution of the precedence effect in transmission from co-colonization as a force in coexistence hierarchies between two strains. Indeed, for any value of coinfection prevalence and overall *R*_0_, hence for any *µ*, the relative contribution of the transmission rate differential between strains (Δ*b*) is higher than that of the transmission bias differential from coinfected hosts (Δ*ω*). It is also interesting to notice in the above expression for 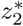 that the relative contribution of the between-strain difference in duration of carriage (Δ*ν*) depends on *µ* but also on the absolute value of the duration of colonization itself *γ*. Essentially, keeping all else fixed, differences in duration of carriage between two strains matter more for their relative fitness, if the colonization episodes are shorter (*γ* higher), and if hosts are longer lived (*r* higher).

Perfectly balancing selection, in this model, under a given *k* ≠ 0, would require that the linear combination of Δ*ν*, Δ*b*, Δ*ω* in the round brackets be equal to zero. Trait variation possibilities satisfying this requirement are infinite and constitute a plane in 3-d in this case, necessarily encoding trade-offs across different fitness dimensions that would as a whole lead to the same epidemiologic fitness for the two strains. Other particular values of 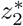 can be studied explicitly and investigated similarly in terms of the constraints implied for the linear combination of Δ*ν*, Δ*b*, Δ*ω*, and other fitness dimensions eventually, if such variation exists.

## 6 Model applications: a roadmap

In earlier theoretical work, interaction among co-infecting agents has been assumed to occur only between different strains, and studied for arbitrary infection multiplicity (Adler and Brunet, 1991). Later evolutionary frameworks, based on (van Baalen and Sabelis, 1995), have considered a full model including same-strain coinfection, but modeling vulnerability to co-infection with a single parameter (Alizon, 2013). This aggregation of within- and between-strain interactions into a net parameter can be found in other co-infection models, considering altered susceptibility to coinfection in the context of disease persistence (Gaivão et al., 2017), and diversity in other traits, e.g. virulence (Alizon et al., 2013) and antibiotic resistance (Hansen and Day, 2014). These studies highlight the importance of coinfection and its epidemiological details for persistence and evolution of microorganisms. Sometimes very complex multi-scale models have been invoked to generate coexistence between strains via coinfection, embedding an explicit within-host dynamics framework (Davies et al., 2019). We argue that many such coinfection and co-colonization models could be mapped to phenomena in the overarching model proposed here, as special cases, or expansions of a particular parameter.

With the here-proposed explicit framework, the impact of coinfection becomes very easy to understand, via the role of the parameter *µ*, given by the ratio of single - to co-infection prevalence in the system, which modulates the relative weight of different trait asymmetries (*θ*_*i*_’s) among strains, and even tuning the net asymmetries in some traits, as is the case for *k*_*ij*_. This role of coinfection (in terms of 2 global determining parameters *R*_0_, *k*) can be studied at a deeper level, at a higher resolution in terms of potential asymmetries within and between strains, and in an entirely analytically-explicit manner which enables precise predictions. These advantages can lead to new applications to study coexistence and vaccination effects in polymorphic systems, going beyond current theoretical insights (Lipsitch, 1997; Gjini et al., 2016). Similarly, our modeling framework could also help obtain clear and direct analytical insights into antibiotic resistance evolution, as an alternative or as a complement to the more cumbersome simulation route (Davies et al., 2019). Below we sketch briefly some ways in which the model can be applied.

### 6.1 Antibiotic treatment, fitness costs and competitive release

We can apply this model to understand epidemiological competition dynamics between two strains under antibiotics, co-circulating in a host population with the possibility of coinfection, which constitutes a big study field in the epidemiological literature (Mulberry et al., 2019). In its simplest form, broad-spectrum antibiotic treatment can be modelled as an increase in global clearance rate of colonization *γ*, keeping fixed all other parameters between strains. Suppose strain 1 is superior in transmission, with strain 2 suffering a relative fitness cost Δ*b >* 0. For variation in duration of carriage we explore both an advantage to strain 1 Δ*ν >* 0 or advantage to strain 2 Δ*ν* < 0. Without perturbation in coinfection parameters, under such scenario, there would be competitive exclusion of the less fit strain, seen earlier in Section 4.1.

But under interactions through coinfection vulnerabilities, coexistence among the two strains is possible. Thus we may explore, the frequency of strain 2, under such scenario, which would then correspond to the variable *z*_2_ in our model. We examine how the equilibrium value of the frequency of strain 2, varies as a function of *γ* (or total broad-spectrum antibiotic treatment), for different values of strain variation in relative duration of carriage (clearance rate) Δ*ν*, and relative transmissibility Δ*b*.

Fixing for example the rescaled co-infection susceptibility matrix to *α*_*ii*_ = *α*_*jj*_ = 0 and 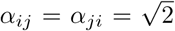, corresponds to the case analyzed earlier, where within-strain susceptibilities to coinfection are lower than between-strain susceptibilities, a condition that *a-priori* favours coexistence. Hence, applying the earlier results, we have that for any (*β, γ*, Δ*b*, Δ*ν*), stable coexistence of two strains is possible only for *T*_1_ (*γ*) ≤ Δ*ν* ≤ *T*_2_ (*γ*). The value of 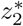 under such condition is given by Eq. 5.9. Thus, in Figure 8, we only consider the values of global clearance rate *γ* guaranteeing coexistence. It is interesting to point out that depending on how the fitness differential between the strain 1 and its competitor strain 2, is manifested (Δ*b*, Δ*ν*), increasing antibiotic administration in a population can have opposing effects: it can increase or decrease the prevalence of a focal strain (see Fig.8a solid vs. dashed lines). This behavior can be understood in full analytic detail because of the explicit expression for strain frequencies, allowing us to compute and verify directly the first and second derivatives of *z*_2_ with respect to *γ* (see Supplementary Material S5).

**Figure 8:**
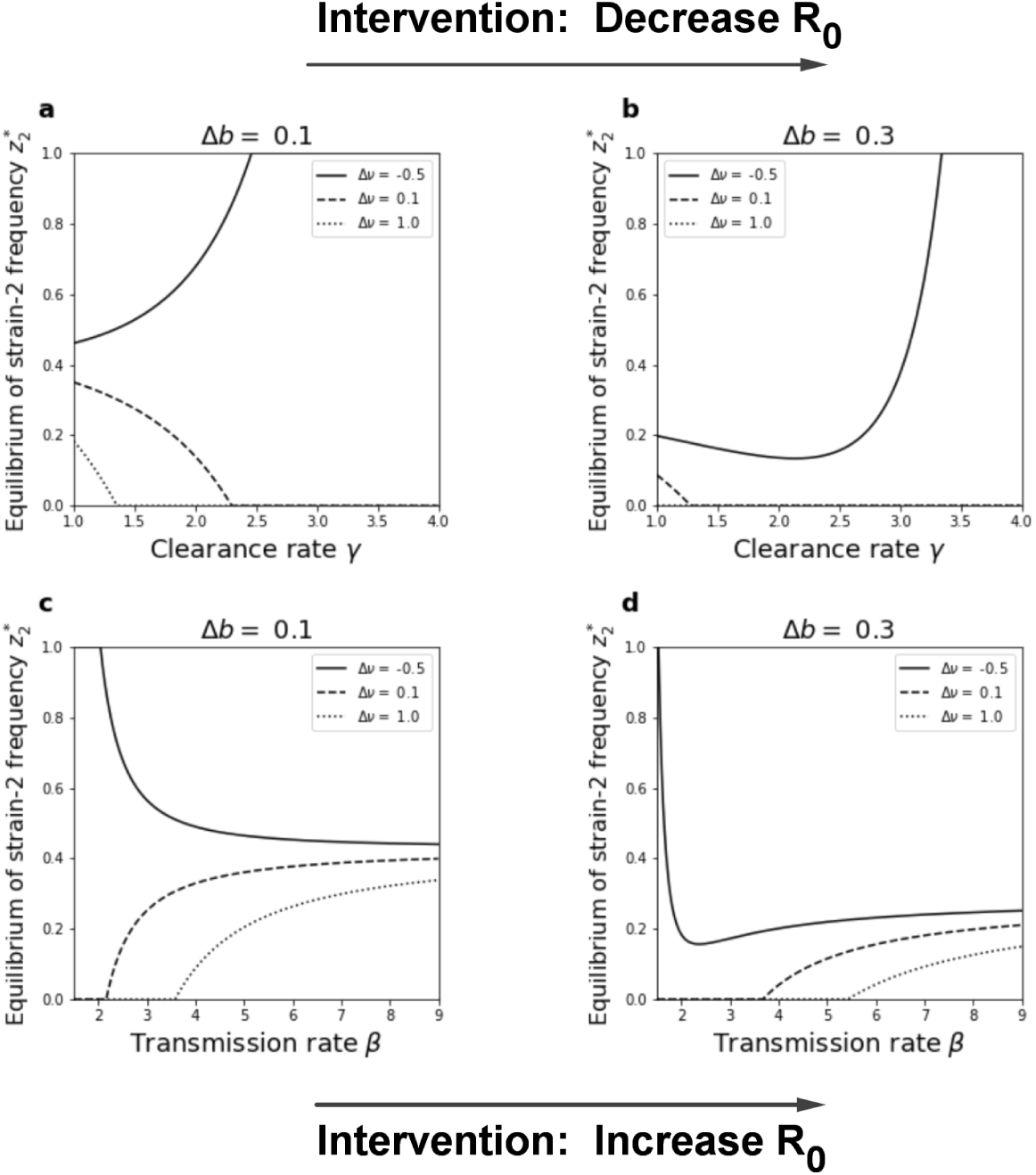
Increasing global clearance or global transmission can have opposite effects on single strain frequency at the endemic equilibrium, depending on underlying trait variation. We plot the frequency of strain 2, 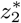 at the coexistence equilibrium (Eq. 5.9), as a function of mean transmission rate and mean clearance rate. Variation among 2-strains is encoded in the transmission and clearance rate axes: *β*_*i*_ and *γ*_*i*_, and co-colonization vulnerabilities *k*_*ij*_. In this simulation, we choose *r* = 0.2, *k* = 1 and the matrix of standardized interactions is assumed symmetric 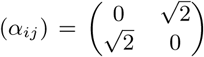, favouring coexistence with *α*_*ij*_ = *α*_*ji*_ *> α*_*ii*_ = *α*_*jj*_. **a-b**. Equilibrium frequency as a function of strain-transcending *γ*. We plot 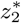 as a function of mean clearance rate *γ* (varied between 1 and 4) for 2 cases of fitness differentials in transmission Δ*b* and 3 cases of variability in clearance Δ*ν*. The global transmission rate is *β* = 4.5 to ensure *R*_0_ *≥* 1. **c-d**. Equilibrium frequency as a function of strain-transcending *β*. We plot 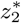, as a function of mean transmission rate *β* (varied between 1.5 and 9) for 2 cases of different variation Δ*b* and 3 cases of Δ*ν*. In these plots, overall clearance rate is held fixed at *γ* = 1 to ensure *R*_0_ *≥* 1.

To understand the additional effects of possible variations in transmission probability from mixed coinfection between strains, we have repeated the same simulations with Δ*w* < 0, favouring strain 2 (Supplementary figure S7). It is clear that also this dimension of fitness (within-host advantage) has a substantial effect on the net competitive dynamics between the two strains, and in particular, in this case, enhances the possibility of two-strain coexistence.

### 6.2 Vaccination, coexistence and strain replacement in colonizing bacteria

Similarly, universal vaccination that protects against both strains could be modelled, to a first-order approximation, as a global reduction of *β* in the system, realized via reduced average susceptibility of all hosts to infection. In figures 8c and 8d, we explore the effect of a universal reduction in *β* on the relative prevalence of two strains. As shown earlier, coexistence is possible only for *S*_1_ (*β*) ≤ Δ*b* ≤ *S*_2_ (*β*). The value of 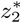 under such condition is given by Equation 5.9. Plotting this as a function of *β* in Figures 8c and 8d, we observe again that changes in strain-transcending transmissibility can have opposing effects on prevalence of a focal strain (here strain 2). They can either favour its increase in prevalence or its decrease, depending on the underlying basic trait variation (Δ*b*, Δ*ν*), as well as on coinfection parameters (*k*_*ij*_ or others).

Turning the inequalities around, another way to interpret the critical borders for Δ*b* is with regards to a strain-specific vaccine. For example, assuming universal coverage, in order to predict the minimal vaccine efficacy needed to exclude a given strain (or group of strains) from the system, given everything else fixed at pre-vaccine baseline, we can use the model to extract the Δ*b* that violates the coexistence inequality criterion: *S*_1_ (*β*) ≤ Δ*b* ≤ *S*_2_ (*β*). Notice that this criterion specifically implies that in populations with different overall *β* a different Δ*b* (targeted vaccine efficacy) may be needed to reach exclusion. In the case of Δ*b* < *S*_1_(*β*) we would ensure exclusion of strain 1 from the system, whereas making Δ*b > S*_2_(*β*) would shift the system to the exclusion of strain 2 steady state.

### 6.3 Effects of host demography on 2-strain epidemiological competition

Changes in natural susceptible recruitment rate (equal host mortality rate) *r* affect *R*_0_ in the system, which also determines *µ*, besides appearing itself also explicitly in Eq.5.10. So even with everything else fixed (*β, γ, k* and Δ*b*, Δ*ν*) it is possible to influence competitive dynamics between strain 1 and strain 2, just via host demography. Increasing host mortality rate, decreases *R*_0_ in the system which increases *µ* and hence gives a larger weight to the trait variation in clearance rate of infection Δ*ν*. Increasing host turnover rate might then enable coexistence of two strains (maintenance of the less fit strain e.g. strain 2) because it could amplify, and even overturn, the relative advantage in duration of carriage (Δ*ν* < 0) versus the disadvantage in transmissibility (Δ*b >* 0). This would imply that in different populations, with different rates of susceptible host turnover, the dynamics of the same two strains could be different. Thanks to this model, all these mechanisms and special cases in competitive dynamics between two closely-related strains at the epidemiological scale can be studied in a fully parameter-explicit and analytical manner, which should promote easier and more direct testable links with data.

### 6.4 Dynamical transitions: from *N* = 2 to the *N* -strain ecological network

Throughout this study, we have shown explicitly and illustrated in detail how global parameters of the neutral model, embedded in the center of the dynamics, can shift qualitatively and quantitatively the net competitive outcome for any given pair of strains. Recall that the *N* = 2 system forms the basic unit in the full competitive network among an arbitrary number *N* of strains, and that qualitative shifts in each network ‘edge’, as a function of global parameters, can have far-reaching effects on the collective dynamics among multiple strains, even when strains differ just in co-infection vulnerabilities (Gjini and Madec, 2021a). Having exposed new and nonlinear gradient effects of *µ, R*_0_, *k, γ, r* in the more complete 2-strain system with variation along 5 fitness dimensions, opens the way towards deeper analysis of their higher-level effects on the *N* − strain assembly, dynamics, and coexistence (Le et al., 2021). Studying dynamical transitions mediated via coinfection prevalence and strain-transcending epidemiological parameters, as well as the statistical distributions of trait variations, possibly informed by data (Abdullahi et al., 2012; Cobey and Lipsitch, 2012; Davies et al., 2019) in the full system, is the natural and exciting next step.

## 7 Discussion

Coinfection is an important aspect of many infectious diseases, and a metaphor applied to model also certain information propagation and species colonization processes. There are substantial modeling efforts dedicated to co-infection in the last decades (Adler and Brunet, 1991; Lipsitch, 1997; Gjini et al., 2016; Alizon et al., 2013; Hansen and Day, 2014; Davies et al., 2019). Yet, simple and sufficiently general mathematical frameworks to analyze and unify the full spectrum of hierarchical patterns emerging from co-infection interactions and variation in other fitness dimensions between two strains are missing. Here, we contribute to fill this gap, thanks to a model reduction obtained after assuming strain similarity (Le et al., 2021). Focusing on *N* = 2, here we have modeled simultaneously 5 fitness dimensions where two strains can differ, and used the decomoposition into two timescales to simplify their dynamics: neutral dynamics between types on a fast timescale and non-neutral selective processes on a slow timescale, driven explicitly by trait variation, going beyond (Gjini and Madec, 2017) where only pairwise vulnerabilities to co-infection (a single ‘trait’) were studied.

Many studies of coinfection are either totally epidemiological in nature (Martcheva, 2009; Thieme, 2007; Lipsitch, 1997), exploring transmission dynamics of infectious agents in a host population, or they focus on evolution of specific pathogen traits (often virulence) (Mosquera and Adler, 1998; Alizon et al., 2013) using the coinfection framework developed by (van Baalen and Sabelis, 1995) for microparasites causing persistent infections. The latter group of studies, typically derive the conditions of invasion of a rare mutant in a host population already infected by a resident strain, following adaptive dynamics theory (Meszéna et al., 2005), where it is further assumed that the resident strain is at equilibrium, that is, that the densities of susceptible, singly-infected and coinfected hosts have reached their equilibrium values. Invader fitness is then evaluated using the basic reproduction ratio (van Baalen and Sabelis, 1995), where it becomes clear that the fitness of a mutant strain is the sum of two components: the fitness achieved through the infection of susceptible hosts and the fitness achieved through the infection of hosts already infected by the resident. Typically by analyzing whether the reproduction ratio is greater than or lower than 1, conditions for successful or unsuccessful invasion, and ultimate evolutionary dynamics for the trait in consideration are established. However, sometimes the criteria derived in such models can be model-dependent, involve cumbersome mathematical expressions, and may not provide immediate comprehensive insight into the biological mechanisms. Furthermore, explicit frequency dynamics post-invasion are typically not derived or elaborated upon.

In the present work, we have bridged these fields of study. With a rather generic model, we have revealed coexistence and competition mechanisms in their bare essence, and have integrated, generalized and advanced analytically the epidemiological and adaptive dynamics perspectives on coinfection. We have linked population dynamics of endemic transmission with slow selective dynamics in strain trait space, and shown that such dynamics are given by a replicator equation involving the mutual invasion fitness matrix between strains (Eq.2.7). We have generalized single trait evolution to multiple trait evolution, exploring phenotypic differentiation along 5 dimensions between two strains: transmission, clearance, vulnerability to coinfection, duration of coinfection, and transmission biases from mixed coinfection, all of which contribute to mutual invasion fitness (Eq.3.1). We have illustrated the utility of an analytical expression for explicit frequency dynamics between two strains, under an endemic global equilibrium, which allows to use the well-known replicator equation to make predictions for exclusion vs. coexistence (Section 3, Eq.5.1), relative strain abundances over time, the means of different traits, and exposes how the means of different traits act to shape their slow variance dynamics.

By elaborating several systematic examples, we have shown that coinfection effects can be very complex on the epidemiological competition between two strains. Indeed, high coinfection prevalence (small *µ* here) is not always a promoter of coexistence; instead, its effect to generate or prevent polymorphism is non-monotonic, and crucially depends on the type and level of existing phenotypic differentiation between strains. We have also proven formally what type of trait variation is required for 2, 3, and 4 qualitative shifts in the same system as a function of coinfection prevalence alone (Table 3). This may prove extremely relevant when interpreting different ecological outcomes potentially arising between the same strains when they occur together in different geographical locations, environmental settings, temperature, resources or other biophysical gradients that act on *R*_0_ and *k*.

Although we have illustrated special cases, where the traits are uncorrelated, possible co-variation constraints or trade-offs between different traits (e.g. transmission-clearance, or transmission -competition in co-colonization) can be studied under the same analytically-explicit framework, especially for higher *N*, provided they do not violate the similarity assumption. One mathematical requirement for the singular perturbation expansion to hold is small *ϵ*. Yet, numerically we find that the slow-fast approximation remains valid and quantitatively reasonably close to the original system even for values of *ϵ* in the order of 0.2-0.3, expanding its applicability. On the other hand, full characterization of the selective dynamics between two strains for small *ϵ*, as provided by our framework, has utility for studying local bifurcations near neutrality and the origin of speciation in such systems. Inevitably, the feature of expressing multivariate phenotypic differentiation into a common currency, *ϵ*, is central to the slow-fast representation (Le et al., 2021), and restricts somewhat the types of systems that can be modelled to those where multiple traits between strains display similar variance. In practice, there are many ways to define the neutral model for the original system 2.1 starting from given parameters. Although they will lead to slightly different fast-slow dynamics, all of them will eventually be *ϵ* − close to the real dynamics and preserve its key features.

It is important to keep in mind that our model is deterministic in nature, and as such it cannot account for the subtleties in selection dynamics induced by demographic stochasticity, possibly even in cases where net invasion fitnesses fully equalize between the two strains. It has been shown using simpler population models that how net fitness is realized in terms of vital rates between two species can have drastic effects on stochastic extinction or mutual coexistence (Parsons et al., 2008; Lin et al., 2012; Constable et al., 2016). This aspect in our model remains to be studied in the future with new frameworks that incorporate stochasticity. Similarly, extensions in model structure, for example sequential instead of direct clearance from coinfection, are likely to require specific investigation to establish whether a similar replicator-like equation is possible to derive using the singular perturbation approach, and determine the eco-evolutionary feedbacks governing strain selection. The very same techniques used here to explore the role of coinfection can have may potential advantages if applied in the context of more complex host-pathogen systems where host population is structured differently, for example heterogeneity in host-specific traits, or contact networks (Chen et al., 2017; Hébert-Dufresne and Althouse, 2015). Another path for future exploration is clearly linking our theoretical results to the broader theoretical developments in quantitative genetics, also based on Taylor expansions around mean traits, for example the oligomorphic approximation by (Sasaki and Dieckmann, 2011) for examining the joint ecological and evolutionary dynamics of populations with multiple interacting morphs, and recent extensions thereof (Lion et al., 2021).

However, even if arising from a specific epidemiological context, we believe our analytical approach, derived for general number of strains *N* in (Le et al., 2021), starting with a complete coinfection structure, involving both same strain and mixed co-infection as in (van Baalen and Sabelis, 1995; Alizon, 2013; Gjini et al., 2016), and made possible via separation of timescales and model reduction under strain similarity (Gjini and Madec, 2017; Madec a can have countless further theoretical and practical applications, not least by harnessing the elegant simplicity and historical study record of the replicator equation (Hofbauer and Sigmund, 2003). Understanding more deeply and in explicit time-scales multiple-trait selection in systems with co-infecting interacting entities will enable easier and more direct insights on several important health challenges, including antibiotic resistance, virulence evolution, optimal immunization, and patterns of diversity in multi-strain endemic pathogens or colonization-based ecosystems.

## Acknowledgements

Thi Minh Thao Le was supported with a Ph.D. scholarship by the University of Tours. Erida Gjini was supported by PESSOA grant 5666/44637YE awarded by Portuguese Foundation for Science and Technology (FCT), and cooperative agreement 2020-2001-230-Y20V7 as Le Studium Visiting Researcher, from Loire Valley Institute of Advanced Studies, Orleans and Tours, France, hosted by Institut Denis Poisson (CNRS/Universite de Tours/Universite d’Orleans) in 2021.

## Code availability

An illustrative code that simulates the dynamics between two strains in our SIS model with coinfection and multi-trait variation, under the proposed slow-fast approximation and replicator equation, is provided on Github.

## Footnotes

1 Thus 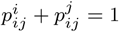 and 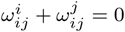.

