## Supplementary Material for "Disentangling how multiple traits drive 2 strain frequencies in SIS dynamics with coinfection"

### Supplementary information

**Disentangling how multiple traits drive 2 strain frequencies in SIS dynamics with coinfection**
**Thi Minh Thao Le** [1,*], **Sten Madec** [1,+], **Erida Gjini** [2,†]
[1] Laboratory of Mathematics, University of Tours, Tours, France,
[2] Center for Stochastic and Computational Mathematics, Instituto Superior Tecnico, Lisbon, Portugal
*, †

### S1 Strain-specific $R_0$ in co-infection vs. actual trait variation

We may consider the relations between the basic reproductive numbers $R_{0,1}$ and $R_{0,2}$ in determining the winner in the case of exclusion. We recall the basic reproductive number of each strain, see (Le et al., 2021) as follows

$$R_{0,1} = \frac{\beta}{m} + \epsilon\frac{\beta}{m}\left(b_1 - \frac{\gamma\nu_1}{m}\right) + O\left(\epsilon^2\right), \qquad R_{0,2} = \frac{\beta}{m} + \epsilon\frac{\beta}{m}\left(b_2 - \frac{\gamma\nu_2}{m}\right) + O\left(\epsilon^2\right) \tag{S1.1}$$

which implies

$$R_{0,1} \geq R_{0,2} \iff b_1 - b_2 \geq \frac{\gamma}{m}\left(\nu_1 - \nu_2\right). \tag{S1.2}$$

Strain 1 has higher strain-specific $R_0$ if and only if its advantage in transmission is bigger than its relative disadvantage in clearance rate, weighted by the global reproduction number.

For any given value $R_{0,1}$, when we fix any value of $\nu_1$, we can find unique value $b_1$ such that strain 1 is associated to $b_1$ and $\nu_1$ and has the basic reproductive number $R_{0,1}$. The same holds true for strain 2. This is plausible because of the formulas $R_{0,i} = \dfrac{\beta_i}{m_i} = \dfrac{\beta\left(1+\epsilon b_i\right)}{m+\epsilon\gamma\nu_i}$.
Hence, with only values of basic reproductive number $R_{0,1}$ and $R_{0,2}$, in a coinfection model, we cannot determine the long time behavior of the dynamics. Figure S1 is actually a vertical slice in each sub figure of figure S6 when we keep $\Delta\nu$ unchanged and change the value of $\Delta b$ to vary $(R_{0,1}, R_{0,2})$. It is consistent with figure S6 when the smaller $\Delta\nu$ leads to the smaller possibility of coexistence in the same range of $\Delta b$ but the possibility of the exclusion of strain 2 stays the same.

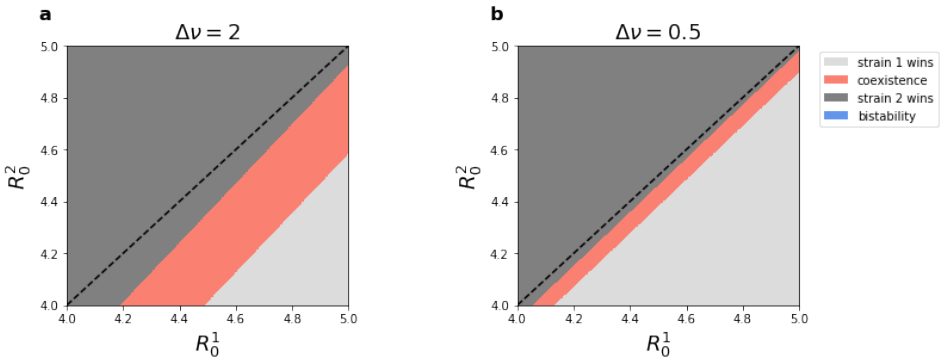


Figure S1: **Ecological scenarios do not depend just on relative basic reproduction numbers of strains $R_{0,1}$ and $R_{0,2}$.** We compare the case of large variation in clearance rate between two strains $\nu_1 = -1$, $\nu_2 = 1$ (**a**) with lower one $\nu_1 = -0.25$, $\nu_2 = 0.25$ (**b**). The other parameters are assumed: $k = 3$, $m = 1.5$, $\beta = 6$, $\epsilon = 0.1$. We vary $\Delta b$ in each case to obtain the relative comparison between $R_{0,1}$ and $R_{0,2}$ shown in the figures. The variations are in transmission rates $\beta_i$, single clearance rates $\gamma_i$, and for coinfection clearance rate we assume $\gamma_{ij}$ with $u_{12} = u_{21} = \min\{\nu_1, \nu_2\}$. Comparing **(a)** and **(b)**, for the same combination of strain-specific $R_0$, the scenarios can diverge depending on the actual difference in strain-specific parameters leading to the particular reproduction number. Particularly, where the $\Delta\nu$ is higher (a), the relative benefit of strain 2 from lower clearance in mixed co-colonization is larger, leading to a larger strain-2 only region, and a larger coexistence region, over the same $R_0$ range. (Code)

### S2 Speed of strain dynamics depends on global parameters

Our model allows explicit quantification of the speed of strain dynamics as a function of epidemiological parameters. Next we illustrate a dynamics example for a 2-strain system tending to exclusion. For the same relative variation between two strains, the dynamics are much faster when $R_0$ is lower, in this case obtained by changes in $\beta$. For the dynamics in figure S2, we calculate the values of theta's and pairwise invasion fitnesses in each sub figure as follows:

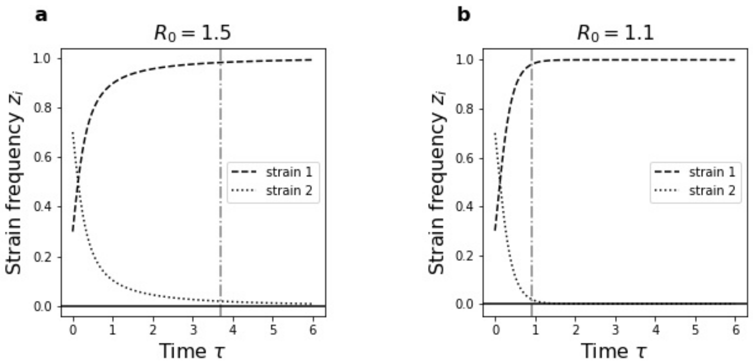


Figure S2: **An example of 2-strain system with multi-trait variation.** $b_1 = 1, b_2 = -1.5$, $\nu_1 = -2, \nu_2 = -3$, $(u_{ij})_{ij} = \begin{pmatrix} 1 & -2 \\ -2 & 3 \end{pmatrix}$, $(\omega^i_{ij})_{ij} = \begin{pmatrix} -1 & -3 \\ -1 & 3 \end{pmatrix}$ and $(\alpha_{ij})_{ij} = \begin{pmatrix} -\sqrt{2} & 0 \\ \sqrt{2} & 0 \end{pmatrix}$. We choose mean transmission rate $\beta = 4$, mortality $r = 0.2$, mean co-colonization interaction factor (altered susceptibility to co-colonization) $k = 1.5$, and the initial values of strain frequencies $(z_1, z_2) = \begin{pmatrix} 0.3 & 0.7 \end{pmatrix}$. Both figures present periods of transient coexistence before the exclusion of strain 1. The dashed-dotted grey vertical lines, denoted by $d$ in two figures show the region where $\min(z_1, z_2) > 0.02$. Figure **(a)** plots for $R_0 = 1.5$, in which, $d$ goes through a point between 3 and 4 (near 4). Figure **(b)** plots for $R_0 = 1.1$, in which, $d$ goes through a point very near and less than 1. Thus, it can be seen that the the region of "effective transient coexistence" when $R_0 = 1.5$ is larger than when $R_0 = 1.1$.

- **(a)**: $\begin{pmatrix} \theta_1 & \theta_2 & \theta_3 & \theta_4 & \theta_5 \end{pmatrix} \approx \begin{pmatrix} 0.48 & 0.2 & 0.09 & 0.03 & 0.2 \end{pmatrix}$, and $\left(\lambda_1^2, \lambda_2^1\right) \approx (10.37, -0.20)$.
- **(b)**: $\begin{pmatrix} \theta_1 & \theta_2 & \theta_3 & \theta_4 & \theta_5 \end{pmatrix} \approx \begin{pmatrix} 0.51 & 0.39 & 0.03 & 0.07 & 0.0 \end{pmatrix}$, and $\left(\lambda_1^2, \lambda_2^1\right) \approx (8.28, -4.42)$.

It is easy to verify that because of the values of $\lambda_i^j$, we have exclusion of strain 2 from the system and only strain 1 persists, but depending on the value of $R_0$ the dynamics will be quicker or slower. In this particular case, reducing $R_0$, hence overall endemic prevalence of the two strains, leads to faster exclusion dynamics and shorter period of transient coexistence.

### S3 Qualitative transitions in the same system when varying $\mu$

#### S3.1 A result for variation occurring only in two out of: i) transmission rates $\beta_i$, ii) clearance rates $\gamma_i$ and ii) transmission biases from coinfection $p_{ij}^i$

In this subsection, we prove that, in the case of variation between 2 strains only, can be in transmission rates $\beta_i$, clearance rates $\gamma_i$ and transmission probability from co-colonized hosts, there is at most one shift for the ecological outcome as a function of $\mu$ (single to coinfection ratio in the system). Indeed, we show this result in three cases.

**Case 1: Transmission rates $\beta_i$ and clearance rates $\gamma_i$ vary between strains.**

By formulae (4.1) with $\Delta\omega = 0$ we have that mutual invasion fitnesses are:

$$\begin{cases} \lambda_1^2 = \dfrac{\gamma}{2(\gamma + r)} \mu (2\mu + 1) \Delta\nu + (\mu + 1)^2 \Delta b \\ \lambda_2^1 = -\lambda_1^2 \end{cases}. \tag{S3.1}$$

It suffices to show that the following equation can not have two positive roots, when solved for $\mu$

$$\frac{\gamma}{2(\gamma + r)} \mu (2\mu + 1) \Delta\nu + (\mu + 1)^2 \Delta b = 0.$$

Assume the contradiction, which means that it has two positive roots, which can be denoted by $\mu_1$ and $\mu_2$. By Viete's theorem, we have that

$$\mu_1 + \mu_2 = -\frac{\frac{\gamma}{2(\gamma+r)} \Delta\nu + 2\Delta b}{\frac{\gamma}{\gamma+r} \Delta\nu + \Delta b} > 0, \qquad \mu_1 \mu_2 = \frac{\Delta b}{\frac{\gamma}{\gamma+r} \Delta\nu + \Delta b} > 0. \tag{S3.2}$$

Thus, (S3.2) implies $\mu_1 + \mu_2 + \dfrac{3}{2} \mu_1 \mu_2 > 0$, which is equivalent to

$$-\frac{\frac{\gamma}{2(\gamma+r)} \Delta\nu + 2\Delta b}{\frac{\gamma}{\gamma+r} \Delta\nu + \Delta b} + \frac{\frac{3}{2} \Delta b}{\frac{\gamma}{\gamma+r} \Delta\nu + \Delta b} > 0,$$

which is absurd because the left-hand side is equal to $-\frac{1}{2}$.

**Case 2: Clearance rates $\gamma_i$ and transmission probability from coinfected hosts $p_{ij}^i$ vary.**

By formulae (4.1) with $\Delta b = 0$ we have that mutual invasion fitnesses are given by:

$$\begin{cases} \lambda_1^2 = \dfrac{\gamma}{2(\gamma+r)}\mu(2\mu+1)\Delta\nu + (\mu+1)\Delta\omega \\ \lambda_2^1 = -\lambda_1^2 \end{cases}. \tag{S3.3}$$

It suffices to show that the following equation can not have two positive roots, when solved for $\mu$

$$\frac{\gamma}{2(\gamma+r)}\mu(2\mu+1)\Delta\nu + (\mu+1)\Delta\omega = 0.$$

Assume the contradiction, which means that it has two positive roots, which can be denoted by $\mu_1$ and $\mu_2$. By Viete's theorem, we have that

$$\mu_1 + \mu_2 = -\frac{\frac{\gamma}{2(\gamma+r)}\Delta\nu + \Delta\omega}{\frac{\gamma}{\gamma+r}\Delta\nu} > 0, \qquad \mu_1\mu_2 = \frac{\Delta\omega}{\frac{\gamma}{\gamma+r}\Delta\nu} > 0. \tag{S3.4}$$

Thus, (S3.4) implies $\mu_1 + \mu_2 + \mu_1\mu_2 > 0$, which is equivalent to

$$-\frac{\frac{\gamma}{2(\gamma+r)}\Delta\nu + \Delta\omega}{\frac{\gamma}{\gamma+r}\Delta\nu} + \frac{\Delta\omega}{\frac{\gamma}{\gamma+r}\Delta\nu} > 0,$$

which is absurd because the left-hand side is equal to $-\frac{1}{2}$.

**Case 3: Transmission probabilities from coinfected hosts $p_{ij}^i$ and transmission rates $\beta_i$ vary.**

By formulae (4.1) with $\Delta\nu = 0$ we have that

$$\begin{cases} \lambda_1^2 = (\mu+1)^2\Delta b + (\mu+1)\Delta\omega \\ \lambda_2^1 = -\lambda_1^2 \end{cases}. \tag{S3.5}$$

The quadratic equation

$$(\mu+1)^2\Delta b + (\mu+1)\Delta\omega = 0$$

has one root to be $-1$. Hence, it can not have two positive roots, which implies our requirement.

#### S3.2 The case of four ecological outcomes in the same system according to $\mu$

It is possible to have four distinct ecological outcomes between which the same system with two strains can shift, as a function of global $\mu$. Initially, we will prove that the necessary condition for the presenting of fully four survival outcomes: E1, E2, C, B is that, variations are in coinfection clearance rates $\gamma_{ij}$ and co-colonization interaction $k_{ij}$.

*Proof.* • **Firstly, we prove that if variations are only in $\beta_i$, $\gamma_i$, $p_{ij}^i$ and $k_{ij}$ then we can not have four survival scenarios as $\mu \to \infty$.**
The formulas for pairwise invasion fitnesses are

$$\begin{aligned} \lambda_1^2 =& \frac{\gamma}{2(\gamma+r)}\mu(2\mu+1)\Delta\nu + (\mu+1)\Delta\omega + \frac{(R_0-1)\mu}{2}(\mu(\alpha_{21}-\alpha_{12}) + \alpha_{21} - \alpha_{22}) + (\mu+1)^2\Delta b \\ \lambda_2^1 =& -\frac{\gamma}{2(\gamma+r)}\mu(2\mu+1)\Delta\nu - (\mu+1)\Delta\omega + \frac{(R_0-1)\mu}{2}(-\mu(\alpha_{21}-\alpha_{12}) + \alpha_{12} - \alpha_{11}) - (\mu+1)^2\Delta b \end{aligned}. \tag{S3.6}$$

We have that two equations $\lambda_1^2 = 0$ and $\lambda_2^1 = 0$ are respectively equivalent to

$$(\mathbf{Eq1}): \quad A_1\mu^2 + B_1\mu^2 + C_1 = 0, \qquad A_1 > 0$$

and

$$(\mathbf{Eq2}): \quad A_2\mu^2 + B_2\mu^2 + C_2 = 0, \qquad A_2 > 0,$$

in which $A_1 = A_2$ and $C_1 = C_2$, easily deduced from (S3.6).

On the other side, by direct verification, to have fully four outcomes, the signs of $\left(\lambda_1^2, \lambda_2^1\right)$ must change at least three times, so one in two equations **(Eq 1)** or **(Eq 2)** must have two distinguished positive solutions and the other must have at least one positive solution. By Viete's theorem, the products of solutions in two equations **(Eq 1)** and **(Eq 2)** are equal. Hence, **(Eq 1)** and **(Eq 2)** must have both positive solutions. Denote two positive solutions of **(Eq 1)** are $x_1 < x_2$, two positive solutions of **(Eq 2)** are $y_1 < y_2$.
By direct checking, and note that when $\mu \to +\infty$ or $\mu \to -\infty$ , $\lambda_1^2 + \lambda_2^1 \to 0$, we have that, to have fully four outcomes, it must be

$$x_1 < y_1 < x_2 < y_2$$

or

$$y_1 < x_1 < y_2 < x_2.$$

Both of these inequality requires the products of solutions in **(Eq 1)** is strictly less or more than the products of solutions in **(Eq 2)**, which is a contradiction of the equality of two products mentioned before.
Thus, if variations are only in $\beta_i$, $\gamma_i$, $p_{ij}^i$ and $k_{ij}$ then we can not have four survival scenarios as $\mu \to \infty$.

• **Secondly, we prove that if variations are only in $\beta_i$, $\gamma_i$, $\gamma_{ij}$ and $p_{ij}^i$ then we can not have four survival scenarios as $\mu \to \infty$.**

The formulas for pairwise invasion fitnesses now read

$$\begin{aligned} \lambda_1^2 =& \frac{\gamma}{2(\gamma+r)}\mu(2\mu+1)\Delta\nu + \frac{\gamma}{2(\gamma+r)}(\mu+1)\Delta_2 u + (\mu+1)\Delta\omega + (\mu+1)^2\Delta b \\ \lambda_2^1 =& -\frac{\gamma}{2(\gamma+r)}\mu(2\mu+1)\Delta\nu + \frac{\gamma}{2(\gamma+r)}(\mu+1)\Delta_1 u - (\mu+1)\Delta\omega - (\mu+1)^2\Delta b \end{aligned}. \tag{S3.7}$$

For the sake of simplicity, we denote that $m := \frac{2(\gamma+r)}{\gamma}$, $d_1 := -u_{12} - u_{21} + 2u_{22}$ and $-d_2 := -u_{12} - u_{21} + 2u_{11}$.
We have that two equations $\lambda_1^2 = 0$ and $\lambda_2^1 = 0$ are respectively equivalent to

$$(\mathbf{Eq3}): \quad A_1'\mu^2 + B_1'\mu^2 + C_1' = 0, \qquad A_1' > 0$$

and

$$(\mathbf{Eq4}): \quad A_2'\mu^2 + B_2'\mu^2 + C_2' = 0, \qquad A_2' > 0,$$

now in which

$$A_1' = 2\Delta\nu + m\Delta b, \qquad B_1' = \Delta\nu + d_1 + 2m\Delta b, \qquad C_1' = d_1 + m\Delta b$$

and

$$A_2' = 2\Delta\nu + m\Delta b, \qquad B_2' = \Delta\nu + d_2 + 2m\Delta b, \qquad C_2' = d_2 + m\Delta b.$$

Denote two solutions of **(Eq 3)** to be $x_3 < x_4$, and two solutions of **(Eq 4)** to be $y_3 < y_4$. By the same arguments, it can be deduce that, to have fully four survival outcome, it must be

$$(*) \qquad x_3 < y_3 < x_4 < y_4$$

or

$$(**) \qquad y_3 < x_3 < y_4 < x_4$$

where at least $0 < x_3 < x_4$ or $0 < y_3 < y_4$. Without loss of generality, similarly to the previous arguments, we assume that $0 < x_3 < x_4$ and $y_4 > 0$.

According to Viete's theorem, we obtain, note that $A_1' = A_2'$

$$x_3 + x_4 = -\frac{B_1'}{A_1'}, \qquad x_3 x_4 = \frac{C_1'}{A_1'},$$

$$y_3 + y_4 = -\frac{B_2'}{A_2'}, \qquad y_3 y_4 = \frac{C_2'}{A_2'}.$$

**Case 1:** $A_1' = A_2' > 0$.

- If $d_1 > d_2$ then $-\frac{B_1'}{A_1'} < -\frac{B_2'}{A_2'}$ and $\frac{C_1'}{A_1'} > \frac{C_2'}{A_2'}$, which implies $x_3 + x_4 < y_3 + y_4$ and $x_3 x_4 > y_3 y_4$. From $x_3 + x_4 < y_3 + y_4$, then $0 < x_3 < y_3 < x_4 < y_4$, which contradicts $x_3 x_4 > y_3 y_4$.

- If $d_1 < d_2$ then $-\frac{B_1'}{A_1'} > -\frac{B_2'}{A_2'}$ and $\frac{C_1'}{A_1'} < \frac{C_2'}{A_2'}$, which implies $x_3 + x_4 > y_3 + y_4$ and $x_3x_4 < y_3y_4$. From $x_3x_4 < y_3y_4$, then $0 < x_3 < y_3 < x_4 < y_4$, which contradicts $x_3 + x_4 > y_3 + y_4$.

**Case 2:** $A_1' = A_2' < 0$.

- If $d_1 > d_2$ then $-\frac{B_1'}{A_1'} > -\frac{B_2'}{A_2'}$ and $\frac{C_1'}{A_1'} < \frac{C_2'}{A_2'}$, which implies $x_3 + x_4 > y_3 + y_4$ and $x_3x_4 < y_3y_4$. From $x_3x_4 < y_3y_4$, then $0 < x_3 < y_3 < x_4 < y_4$, which contradicts $x_3 + x_4 > y_3 + y_4$.
- If $d_1 < d_2$ then $-\frac{B_1'}{A_1'} < -\frac{B_2'}{A_2'}$ and $\frac{C_1'}{A_1'} > \frac{C_2'}{A_2'}$, which implies $x_3 + x_4 < y_3 + y_4$ and $x_3x_4 > y_3y_4$. From $x_3 + x_4 < y_3 + y_4$, then $0 < x_3 < y_3 < x_4 < y_4$, which contradicts $x_3x_4 > y_3y_4$.

Hence, if variations are only in $\beta_i$, $\gamma_i$, $\gamma_{ij}$ and $p_{ij}^i$ then we can not have four survival scenarios as $\mu \to \infty$. □

In the main text, we give an example, in which varying $\mu$ from 0 to $\infty$ may give us 4 outcome exclusion of either strain, coexistence and bistability.

### S4 Examples for 3 possible global outcomes

We illustrate possible outcomes as a function of $\Delta b$ and $\Delta\nu$, similarly to Figure 6, when the perturbations are only in transmission rates $\beta_i$, clearance rates $\gamma_i$ and co-colonization interactions $k_{ij}$. We recall that the borders separating exclusion regions are lines representing $\lambda_1^2 = 0$ and $\lambda_2^1 = 0$.

According to the explicit formulas of $\left(\lambda_1^2, \lambda_2^1\right)$, the border lines have the same slope, thus leading to parallelism. Thus, there are at most 3 possible outcomes for each fixed value $(R_0, k)$. Figures S3 and S4 shows that changing the matrix $(\alpha_{ij})$ may generate different final outcomes.

According to (3.1), we can deduce that regardless of changing $(\alpha_{ij})$, we can not observe coexistence and bistability for a fixed value of $(\alpha_{ij})$. Indeed, according (3.1), two coefficients of $\Delta\nu$ in the formulae of $\lambda_1^2$ and $\lambda_2^1$ are of opposite signs, and the same holds for $\Delta b$.

Then, for $(\Delta\nu, \Delta b) \to (\infty, \infty)$ and $(-\infty, -\infty)$, we have the opposing signs of $\left(\lambda_1^2, \lambda_2^1\right)$, leading to exclusion.

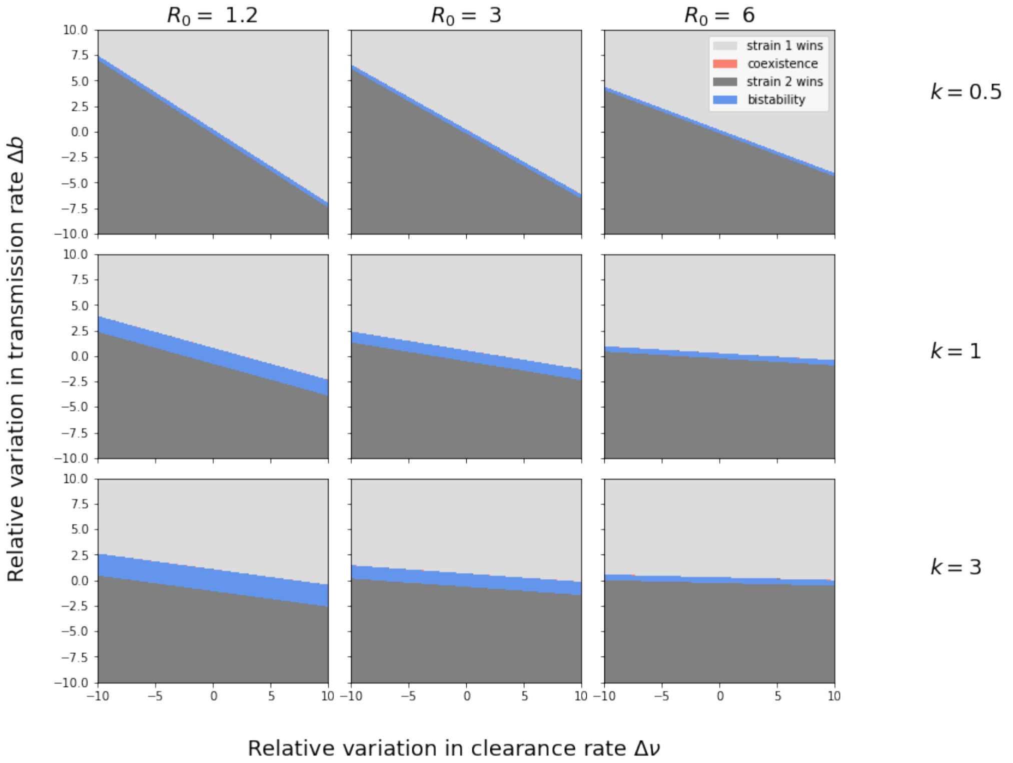


Figure S3: **Illustration of 3 possible outcomes, as a function of $\Delta\nu$ and $\Delta b$, for different values of $k$ and $R_0$.** We highlight the respective regions in different colors, according to the critical relationship between $\Delta b$, $\Delta\nu$, $k$ and $R_0$ when perturbations happen only $\beta_i$, $\gamma_i$, and $k_{ij}$ with $\gamma = 1$, $r = 0.2$ and the matrix $(\alpha_{ij})$ be $\begin{pmatrix} 0 & -\sqrt{2} \\ -\sqrt{2} & 0 \end{pmatrix}$. Three possible survival outcomes include the exclusion of strain 1, bistability and the exclusion of strain 2.

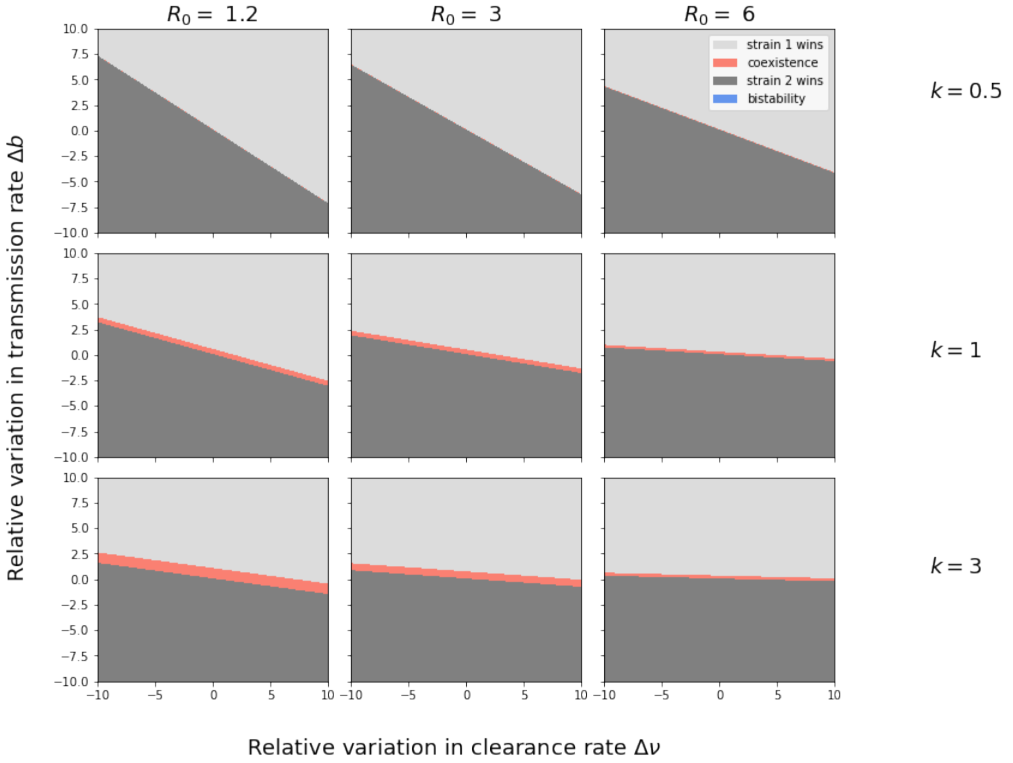


Figure S4: **Illustration of 3 possible outcomes, as a function of difference in transmission and clearance rates between two strains, for different values of $k$ and $R_0$.** We highlight the respective regions in different colors, according to the critical relationship between $\Delta b$, $\Delta \nu$, $k$ and $R_0$ when perturbations happen only $\beta_i$, $\gamma_i$, and $k_{ij}$ with $\gamma = 1$, $r = 0.2$ and the matrix $(\alpha_{ij})$ be $\begin{pmatrix} -2 & 0 \\ 0 & 0 \end{pmatrix}$. This changing of matrix $(\alpha_{ij})$ flips the position of two lines $\lambda_1^2 = 0$ and $\lambda_2^1 = 0$ leads to the other possible outcomes. Three possible system outcomes include: the exclusion of strain 1, coexistence and the exclusion of strain 2.

### S5 The analytical expression for coexistence prevalence

Here we compute the value of $z_2^*$ in the case of symmetric co-colonization interaction matrix. Thus: $\alpha_{ij} = \alpha_{ji}$, $\alpha_{ii} = \alpha_{jj}$. We also assume that strain-specific transmission probability from mixed coinfected hosts is in this case $p_{ij}^i = \frac{1}{2}$ (no priority effects).

- Perturbation in $\beta_i, \gamma_i, \gamma_{ij}, k_{ij}$

$$z_2^* = \frac{1}{2} \cdot \frac{-\Delta b - \dfrac{\theta_2}{\theta_1}\Delta\nu + \dfrac{\theta_3}{\theta_1}\left(2u_{11} - u_{12} - u_{21}\right) + \dfrac{\theta_5}{\theta_1}\left(\alpha_{12} - \alpha_{11}\right)}{\dfrac{\theta_3}{\theta_1}\left(u_{11} + u_{22} - u_{12} - u_{21}\right) + \dfrac{\theta_5}{\theta_1}\left(\alpha_{12} - \alpha_{11}\right)} \tag{S5.1}$$

- Perturbation in $\beta_i, \gamma_i, k_{ij}$

$$\begin{aligned} z_2^* &= \frac{1}{2} \cdot \frac{-\Delta b - \dfrac{\theta_2}{\theta_1}\Delta\nu + \dfrac{\theta_5}{\theta_1}\left(\alpha_{12} - \alpha_{11}\right)}{\dfrac{\theta_5}{\theta_1}\left(\alpha_{12} - \alpha_{11}\right)} = \frac{1}{2}\left(\frac{-\Delta b - \dfrac{\theta_2}{\theta_1}\Delta\nu}{\dfrac{\theta_5}{\theta_1}\left(\alpha_{12} - \alpha_{11}\right)} + 1\right) \\ &= \frac{1}{2}\left(-\frac{1}{\alpha_{12} - \alpha_{11}} \cdot \frac{2\left(\gamma + r\right) + k\left(\beta - \gamma - r\right)}{k\left(\beta - \gamma - r\right)^2} \cdot \gamma\Delta\nu - \frac{1}{\alpha_{12} - \alpha_{11}} \cdot \frac{2\left(\gamma + r + k\left(\beta - \gamma - r\right)\right)^2}{k\left(\beta - \gamma - r\right)^2}\Delta b + 1\right). \end{aligned} \tag{S5.2}$$

#### S5.1 Studying the monotonicity of $z_2^*$: increasing vs. decreasing equilibrium strain frequency

For the sake of simplicity, and for the purposes of illustration, we consider the equilibrium of resistance strain in section 4. Recall the formula of $z_2^*$ in (5.9)

$$z_2^* = \frac{1}{2} - \Delta\nu \cdot \gamma \frac{2(\gamma + r) + k(\beta - \gamma - r)}{2\sqrt{2}k(\beta - \gamma - r)^2} - \Delta b \frac{[\gamma + r + k(\beta - \gamma - r)]^2}{\sqrt{2}k(\beta - \gamma - r)^2}. \tag{S5.3}$$

To investigate the monotonicity of strain frequency, we need to compute the first partial derivative of $z_2^*$ with respect to $\beta$ and $\gamma$, noting that $m = \gamma + r$ and investigate its sign:

$$\frac{\partial z_2^*}{\partial \beta} = -\Delta\nu \frac{1}{2\sqrt{2}k} \frac{(k-4)m - \beta k}{(\beta - m)^3} - \Delta b \frac{1}{\sqrt{2}k} \frac{2m((k-1)m - \beta k)}{(\beta - m)^3}, \tag{S5.4}$$

and

$$\frac{\partial z_2^*}{\partial \gamma} = -\Delta\nu \frac{1}{2\sqrt{2}k} \frac{\beta(\beta k - km + 4m) - r((k+2)\beta - (k-2)m)}{(\beta - m)^3} - \Delta b \frac{1}{\sqrt{2}k} \frac{2\beta(\beta k - km + m)}{(\beta - m)^3}. \tag{S5.5}$$

It can be seen that the expressions are entirely explicit but they display nonlinear dependence on many parameters, including strain variation $(\Delta b, \Delta\nu)$, as well as mean parameters $\beta, m$ or coinfection determinants ($k$, etc.). This allows to obtain a full analytic understanding of the coexistence between two strains at the epidemiological level and how their relative hierarchies depend and may shift with overall context.

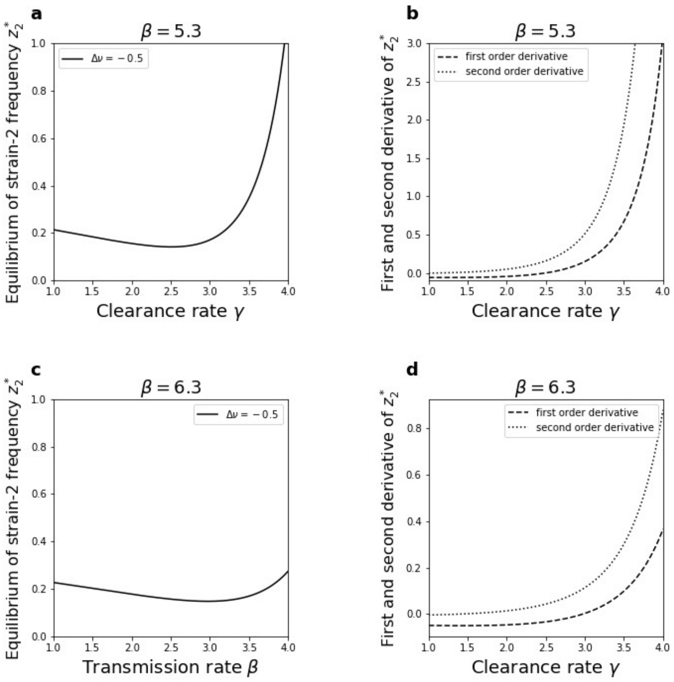


Figure S5: **Monotonicity and convexity of $z_2^*$ according to transmission rate $\beta$ and clearance rate $\gamma$.** We choose $r = 0.2$, $k = 1$ and matrix including entries $\alpha_{ij}$ is $\begin{pmatrix} 0 & \sqrt{2} \\ \sqrt{2} & 0 \end{pmatrix}$. The behavior observed is by plotting for $\Delta\nu = -0.5$. In **(a)** and **(c)**, setting $\Delta b = 0.3$, we plot $z_2^*$ according to $\gamma$ with $\beta = 5.3$ and $\beta = 6.3$ respectively. Figures **(c)** and **(b)** present the curve of first and second orders derivatives of $z_2^*$ according to $\gamma$, respective for figures **(a)** and **(b)**.

#### S5.2 Studying the convexity of $z_2^*$: accelerating vs. decelerating behavior

In this subsection, we consider the convexity of $z_2^*$ with respect to $\beta$ and $\gamma$, global strain-transcending parameters, that can be controlled via interventions (antibiotic treatment, vaccines, etc.). Thus, we compute the second-order derivative of $z_2^*$ related to $\beta$ as follows, noting that $m = \gamma + r$:

$$\frac{\partial^2 z_2^*}{\partial \beta^2} = -\Delta\nu \frac{1}{\sqrt{2}k} \frac{6\beta m + (-4m + k)(\beta - m)}{(\beta - m)^4} - \Delta b \frac{\sqrt{2}m(3m - 2km + 2\beta k)}{k(\beta - m)^4}, \tag{S5.6}$$

and the second-order derivative of $z_2^*$ related to $\gamma$ as follows:

$$\frac{\partial^2 z_2^*}{\partial \gamma^2} = -\Delta\nu \frac{\beta k - (k-6)m}{\sqrt{2}k\,(\beta - m)^4} - \Delta b \frac{2\beta\,((2k+1) - 2\,(k-1)\,m)}{k\,(\beta - m)^4}. \tag{S5.7}$$

These expressions reveal whether the behavior of the strain frequency as a function of global parameters is decelerating or accelerating. This has implications for the control of the strain composition at the population level if we intervene via changing global transmissibility of all strains $\beta$ or global clearance rate $\gamma$. If the behavior is increasing and accelerating, this means that the frequency of that strain can reach fixation for some value of the global parameter. If the behavior is increasing and decelerating (first derivative positive and second derivative negative) this means that the strain frequency saturates at a particular value $< 1$ indicating coexistence in the limit of high values of the control parameter. The following simulation in Figure S5 allows us to see how we can use these analytical expressions above to predict exactly the behavior the strain coexistence frequencies as a function of our parameters, and possible responses to interventions in global transmission rate $\beta$ or clearance rate $\gamma$. In these figures, we can see that $z_2^*$ is always convex related to $\gamma$ in the considered period of varying $\gamma$.

#### S5.3 Effect of transmission priority effects from mixed coinfection $\Delta\omega$

To see the effect of transmission biases from mixed coinfection, as in the section 4 we plot the values of pairwise invasion fitnesses according to the single to co-colonization ratio $\mu$. From 4.1, it is known that if variations are only in transmission rates $\beta_i$, clearance rates $\gamma_i$ and transmission probability from coinfected hosts $p_{ij}^i$, the outcome is always the exclusion of either strain. In this figure, the perturbations are in transmission rates, clearance rates, transmission probability from co-colonized hosts as well as in co-colonization interaction factor $k_{ij}$ to break the anti-symmetry. Except the transmission biases from mixed coinfection $p_{ij}^i$ favor to strain 2, other trait differences in $\beta_i$, $\gamma_i$ favors to strain 1.

We consider in three cases including co-colonization interaction factor $k_{ij}$ favors to strain 2, disfavors to strain 1 and counter balance, respectively. Although the difference $\Delta\omega$ is bias to strain 2, when $\mu$ is large enough, the difference in transmission probability does not effect too much. This can be seen in the similar trending of $\left(\lambda_1^2, \lambda_2^1\right)$ in figures **(a, b, c)** in figure 4, whose values of $\Delta b$ and $\Delta\nu$ are kept the same.

These phenomena are plausible because the weight (written in $\mu$) of transmission probability from coinfected hosts is $\mu + 1$ in the formulae of invasion fitnesses (4.5). Of course, unlike 4.3, when $\mu$ is small enough, $\Delta\omega$ leads to the exclusion of strain 2 in figure S6 **(a)** and the exclusion of strain 1 in figure S6 **(b)**. Meanwhile, $\lambda_1^2$ is always positive in figure 4 **(a)** and $\lambda_2^1$ is always negative in figure 4 **(b)**.

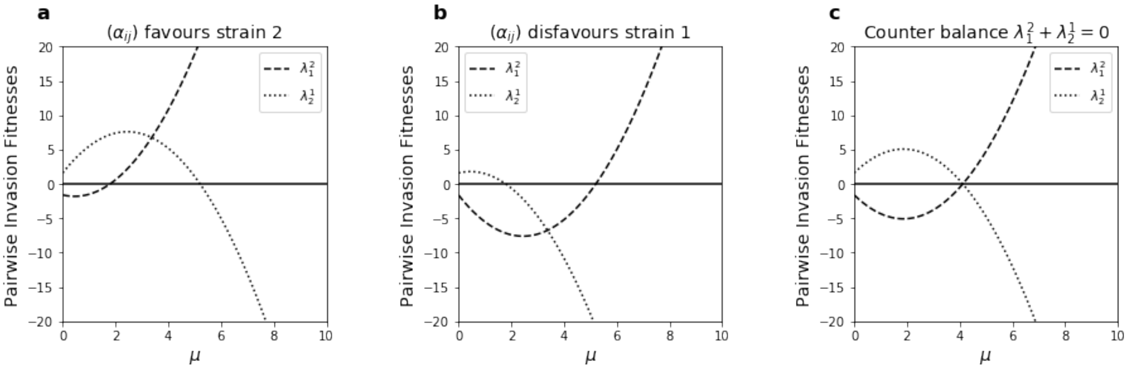


Figure S6: **Additional effects of within-host transmission advantage from mixed coinfection $p_{ij}^i \neq \frac{1}{2}$ (for comparison with Figure 4 a-b-c)** Here we combine variation in co-colonization interactions $k_{ij}$ with variations in transmission rates, infection clearance rates, and transmission probability from coinfected hosts (See Eqs (4.9)). We compute pairwise invasion fitnesses $\left(\lambda_1^2, \lambda_2^1\right)$according to $\mu$ in various cases of co-colonization interaction matrix $(\alpha_{ij})$. We illustrate the cases of transmission and clearance superiority of strain 1 (parameters as in top row of Figure 4): $\Delta b = 0.4$, $\Delta\nu = 0.8$, $R_0 = 5$, $r = 0.5$ and $\gamma = 1.5$. We choose the value $\Delta\omega = -2$ to increase the advantage in transmission probability of strain 2 from mixed coinfection. We consider 3 structures: **(a)** $\begin{pmatrix} -2 & 0 \\ 0 & 0 \end{pmatrix}$; **(b)** $\begin{pmatrix} 0 & 0 \\ 0 & 2 \end{pmatrix}$; **(c)** $\begin{pmatrix} -\sqrt{2} & 0 \\ 0 & \sqrt{2} \end{pmatrix}$ for variation in co-colonization interactions.

Secondly, as mentioned in Sections 6.1 and 6.2, we point out that depending on how the fitness differential between strain 1 and its competitor strain 2, is manifested $(\Delta b, \Delta\nu)$ increasing strain-transcending clearance or tranmission rates ($\gamma$ or $\beta$) in a population can have opposing effects. Besides other trait parameters which are kept unchanged, we analyze the effect of $\Delta\omega < 0$ favouring strain 2, and how this modifies the range of $\gamma$ guaranteeing the coexistence, originally observed in Figure S7. We can see that the shapes of curves representing $z_2^*$ are similar to the Figures 8 **(a, b, c, d)**. respectively. However, in Figures S7 for all **(a, b, c, d)**, for $\Delta\nu > 0$ (which favours strain 1), compared to figures 8 **(a, b, c, d)** respectively, the ranges for survival of strain 2 are larger, highlighting the positive effect of its precedence in transmission from mixed coinfection. In contrast to this, when $\Delta\nu < 0$, the range for coexistence decreases, to a larger advantage of strain 2-only persistence. In all cases, the values of equilibrium of strain 2 in Figure S7 corresponding to each parameter $\gamma$ or $\beta$ are higher than in Figure 8.

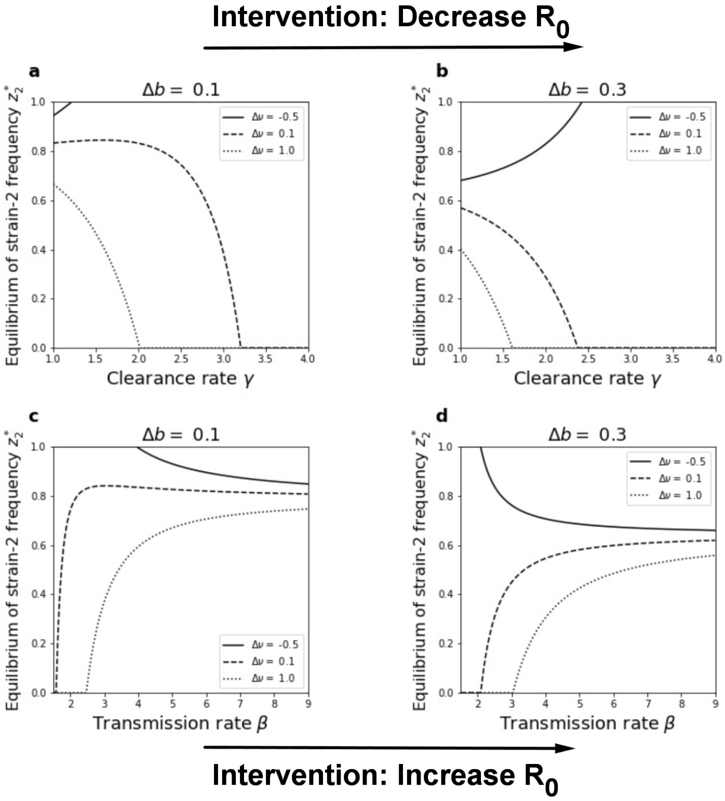


Figure S7: **Strain frequency at the endemic equilibrium vs. $\gamma$ and $\beta$ depends on underlying trait variation (related to Fig.8 but with $\Delta\omega \neq 0$).** We plot the prevalence of strain 2, $z_2^*$ at the coexistence equilibrium (Eq. 5.9), as a function of mean transmission rate and mean clearance rate. Variation among 2-strains is encoded in the transmission and clearance rate axes: $\beta_i$ and $\gamma_i$, transmission probability from co-colonized hosts $p_{ij}^i$ and co-colonization vulnerabilities $k_{ij}$. In this simulation, we keep the same values of $r$, $k$ and the matrix of standardized interactions in Figure 8. We choose $\Delta\omega = -0.5$, which favors strain 2 in precedence of transmission from mixed coinfection. **a-b.** We plot the equilibrium $z_2^*$ as a function of mean clearance rate $\gamma$ (varied between 1 and 4) for 2 cases of fitness differentials in transmission $\Delta b$ and 3 cases of variability in clearance $\Delta\nu$. The global transmission rate is $\beta = 4.5$ to ensure $R_0 \geq 1$. **c-d.** We plot the equilibrium frequency of strain 2, $z_2^*$, as a function of mean transmission rate $\beta$ (varied between 1.5 and 9) for 2 cases of different variation $\Delta b$ and 3 cases of $\Delta\nu$. In these plots, overall clearance rate is held fixed at $\gamma = 1$ to ensure $R_0 \geq 1$.
